# Stability of a biomembrane tube covered with proteins

**DOI:** 10.1101/2022.09.29.510025

**Authors:** Mathijs Janssen, Susanne Liese, Sami C. Al-Izzi, Andreas Carlson

## Abstract

Membrane tubes are essential structural features in cells that facilitate biomaterial transport and inter- and intracellular signalling. The shape of these tubes can be regulated by the proteins that surround and adhere to them. We study the stability of a biomembrane tube coated with proteins by combining linear stability analysis, out-of-equilibrium hydrodynamic calculations, and numerical solutions of a Helfrich-like membrane model. Our analysis demonstrates that both long and short-wavelength perturbations can destabilise the tubes. Numerical simulations confirm the derived linear stability criteria and yield the nonlinearly-perturbed vesicle shapes. Our study highlights the interplay between membrane shape and protein density, where the shape instability concurs with a redistribution of proteins into a banded pattern.

## I. INTRODUCTION

Ubiquitous and essential in biology, phospholipid membranes are self-assembled fluid bilayers of nanometer thickness that compartmentalise biomaterial and form the boundaries of organelles. These bilayer membranes behave tangentially as incompressible fluids in which particles such as proteins can freely diffuse but display elastic behaviour upon deformations in the normal direction. This combination of resisting bend while allowing for tangential flow is a property that leads to a wealth of interesting vesicle shapes on mesoscopic length scales [1]. Of particular interest to biologists and physicists are cylindrical vesicles or membrane tubes, which can be formed from the action of a localised force on a flat membrane. In the simplest case, their radius 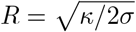 is set by a balance between the bending rigidity of the membrane, *κ*, and surface tension, *σ*[2, 3]. In cells, membrane tubes can be formed by the action of molecular motors [4] and are part of organelles that enable biomaterial exchange and intra- and extra-cellular communication [5, 6]. While some membrane tubes arise transiently [7, 8], others can be long-lived parts of organelles, such as in the peripheral endoplasmic reticulum. In either case, the interaction between the tube membrane and the surrounding proteins is crucial for its emergence, shaping, and stability [9].

As the bilayer is thin compared to the size of the vesicles, it is often modelled as a 2D surface. The classical model for this surface is due to Helfrich, who expressed the free energy of a membrane in terms of its mean and Gaussian curvatures, the lowest order expansion of a general bending geometric free energy in terms of its principle curvatures [10–13]. Zhong-Can and Helfrich showed that a membrane tube of radius *R* becomes unstable if the membrane’s induced curvature is smaller than −1/(2*R*) [14, 15]. This instability, reminiscent of the Rayleigh-Plateau instability of liquid jets, causes membrane tubes to “pearl” for membrane surface tensions *σ* exceeding *σ* > 3*κ*/2*R*^2^ [16–20].

There is extensive literature on how particles—nearby, attached, or inserted into a membrane—affect membrane shape. Examples include the effects of different types of lipids [21–30] polymers [31–35] nanoparticles [36–40] phospholipid-surfactant mixtures [41], protein pumps [20, 42], adsorbed BAR proteins [43–48] inter-calated curvature-inducing proteins [49–51] and crowds of sterically-repelling proteins [52–56] (see also reviews in [57–59] Generally speaking, the total free energy of such systems contains, besides the Helfrich free energy, the free energy of the particles and a term accounting for the interaction of the membrane with the particles. Expressions for the free energy of the particles often contain terms penalising phase boundaries [22, 41, 43, 49, 60] and terms accounting for interactions between the particles and for entropy, either through a Flory-Huggins theory for a mixture of occupied and unoccupied sites [21, 27, 33, 41, 43], or a Ginzburg-Landau expansion thereof [22, 49, 61]. To describe the interaction of particles with a vesicle, most authors [21, 22, 33, 35, 41, 43, 46, 49–51, 60, 62, 63] considered a linear coupling *∝* Λ*Hϕ* of the mean curvature *H* to the areal particle density *ϕ* with a coupling parameter Λ. Leibler showed that such interaction between proteins and a flat membrane sheet changes its effective bending rigidity to *κ−*Λ^2^*/a*, with *a* setting the strength of mutual protein interactions [49]. Consequently, the sheet becomes unstable to ruffles for *κ <* Λ^2^*/a*.

Experiments have shown that membrane tubes with anchored polymers undergo a pearling instability similar to the tension-induced pearling of uncoated membrane tubes [33, 56]. The experimentally-observed tube shapes were interpreted through a theoretical model based on a free energy description of surfaces of constant mean curvature. However, a comprehensive analysis of the stability of protein-covered membrane tubes is still lacking. This paper uses thermodynamic and hydrodynamic approaches to describe the mechanics of membrane tubes with curvature-coupled proteins. We show that in contrast to the flat membrane case, the extrinsic curvature of the tube radius gives rise to a bimodal growth rate for sufficiently high values of protein curvature coupling. When hydrodynamic effects are accounted for, the shorter wavelength instability dominates with a growth timescale set by the membrane viscosity. Finally, we show that there is a nonlinear shape that satisfies that shape equation consisting of constant mean curvature-dilute/dense separated regions with an effective line tension between them. Our results give a range of predictions for the shape of protein-covered membrane tubes that have potential relevance to a broad range of cellular processes, such as ER-Golgi transport [7, 8] and mitochondrial fission [64, 65].

## II. EQUILIBRIUM THEORY

We consider a membrane tube and particles interacting with the membrane and among themselves (see Fig. 1). Without loss of generality, we refer to these particles as proteins. We partition the total free energy *F* = *F*_*H*_ +*F*_*ϕ*_ into a bare membrane term, given by the Helfrich free energy,

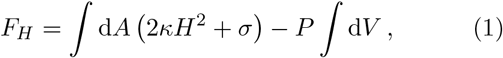

and a contribution *F*_*ϕ*_ due to the proteins. In Eq. (1), *H* is the mean curvature, where our sign convention is such that a cylinder of radius *R* has *H* =−1*/*(2*R*). We omit a Gaussian bending term, which is topologically invariant if we assume the saddle-splay modulus is a constant [12, 66]. In principle, this term could contribute to the energy if the saddle-splay modulus depended on protein concentration. Moreover, *P* is the pressure difference between the fluids inside and outside of the tube, *σ*is the surface tension that acts along the entire membrane, typically around *σ*= 2*×*10^−5^ N m^−1^ [12] and with *κ* = 20*k*_*B*_*T* [67], where *k*_*B*_*T* is the thermal energy.

**Figure 1.**
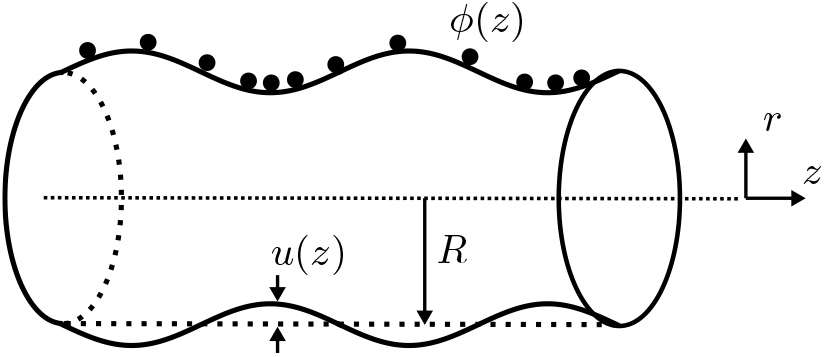
Schematic of a lipid bilayer tube with attached particles.The membrane tube can undergo an instability from a homogeneously coated cylinder of radius R towards an undulating shape *r*(*z*) = *R* + *u*(*z*) and varying areal particle density *ϕ* (*z*). This sketch corresponds to a case for which the protein-membrane interaction parameter Λ > 0. For Λ < 0, bands of large protein concentration are at the crests, instead.

In addition to *F*_*H*_, we consider the following Ginzburg-Landau free energy for proteins and their interaction with the membrane:

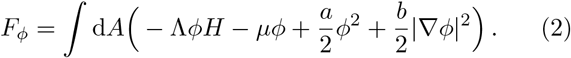

Here, *ϕ* is the areal protein density, satisfying *ϕ≥*0, and ∇ is the surface gradient operator. Λ sets the strength of protein-membrane interactions, *μ* is the energy (per unit area) associated with protein binding to the membrane, *a* sets the protein-protein interactions, and *b* sets the cost of phase separation. We consider *a, b >* 0, such that the last two terms in Eq. (2) favour a vesicle with a homogenous protein coat. By contrast, we consider both Λ *<* 0 and Λ *>* 0. For Λ *>* 0, the first term in Eq. (2) may lead to undulating (“pearling”) tube shapes that minimise the free energy by recruiting proteins (large *ϕ*) to regions where *H >* 0 (the troughs) while depleting them (small *ϕ*) from regions where *H <* 0 (the crests). Conversely, Λ *<* 0 may lead to undulations with increased protein density at the undulating tube’s crests. Experimental data for the values of *a, b*, and Λ are scarce, but we make order-of-magnitude estimates for them in Sec. S1 of the the Electronic Supplementary Information (ESI) [68].

### A. Stability analysis (cylindrical tube)

From hereon, we describe the membrane shape using cylindrical coordinates (*r, θ, z*) and consider a rotationally symmetric tube [20], such that all variables are *θ* independent. We first study the stability of a membrane tube of length *L* and fixed radius *r*(*z*) = *r* with a homogeneous coat of proteins at density *ϕ*(*z*) = *ϕ*. In this case, Eqs. (1) and (2) simplify to

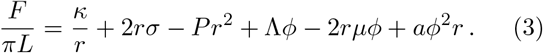

For the free energy *F* to be in a local minimum at a given *r* = *R* and *ϕ* = Φ, three conditions must be met. First, minimising Eq. (3) with respect to the protein density [(*∂F/∂ϕ*)_*R*,Φ_ = 0] yields

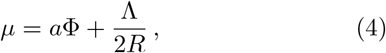

which means that, unlike a membrane sheet [49], a membrane tube has a finite protein density Φ≠ 0 if *μ* = 0. Second, minimising Eq. (3) with respect to the vesicle radius [(*∂F/∂r*)_*R*,Φ_ = 0] yields

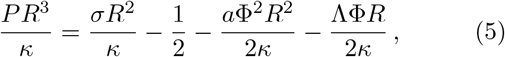

where we used Eq. (4) to eliminate *μ*. While the first two terms on the right-hand side of Eq. (5) represent the conventional Laplace pressure, the last two terms are new and specific to our model problem. Note that by fixing the tube length *L*, we assume to have (indirect) control over the membrane tension *σ*. By contrast, the pressure *P* adjusts itself for given *R, σ*, and other parameters according to Eq. (5).

Finally, the Hessian determinant of *F* must be positive for the tube to be stable. This amounts to [(*∂*^2^*F/∂ϕ*^2^)(*∂*^2^*F/∂r*^2^) − (*∂*^2^*F/∂ϕ∂r*)^2^] _*R*,Φ_ *>*0, giving

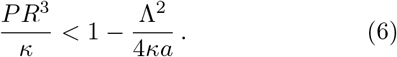

From Eqs. (5) and (6), we conclude that the coated cylinder is stable if

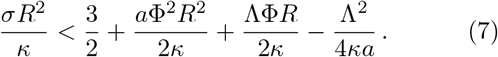

Equation (7) generalises a well-known stability criterion *σR*^2^*/κ <* 3*/*2 for uncoated membrane tubes [11, 18– 20], adding three terms related to the attached proteins. Summarising, for given *σ*, Λ, *μ*, and *a*, a tube of constant radius *R* has a coat of density Φ [Eq. (4)] and a pressure drop *P* [Eq. (5)] across its membrane.

### B. Linear Stability Analysis

To study what happens to the tube if Eq. (7) is not satisfied, we now consider a tube with slight undulations in its radius and coat density, *r*(*z*) = *R*+*u*(*z*) and *ϕ*(*z*) = Φ + *ϕ*(*z*), with *u*(*z*) ≪ *R* and *ϕ*(*z*) *≪* Φ (see Fig. 1). We use that 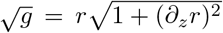, with *g* being the metric determinant, and that [20, 69, 70]

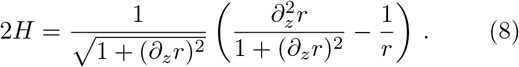

From hereon, we use the following dimensionless variables and parameters: 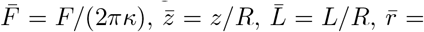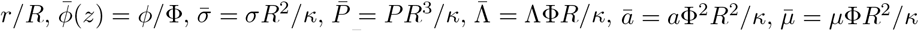 and 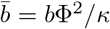. We can then express 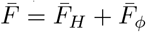 as

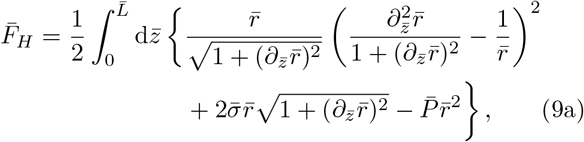

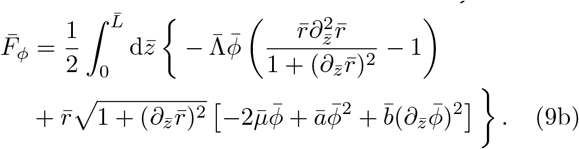

For tubes with radii around *R∼*4 μm, we have 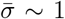. For the other dimensionless parameters, 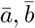, and 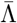, we will consider values around 1 [68].

Next, we expand the perturbations 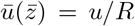 and 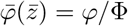 into Fourier modes,

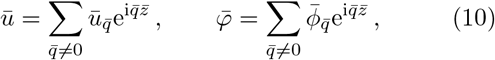

where 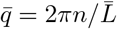 are dimensionless wave numbers (with *n∈*ℕ) and where 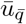 and 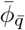 are the amplitudes of the modes. Inserting Eq. (10) and 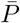 from Eq. (5) into 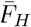 and 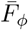 and retaining quadratic terms leads to [68]

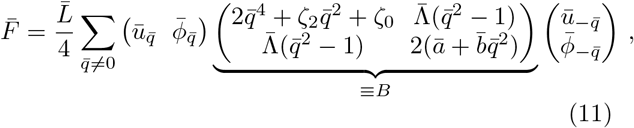

with 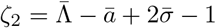 and 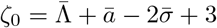. Considering Eq. (11) in the flat membrane limit *R → ∞*, we fou(nd (see Ref. [68]) a leading order contribution at *𝒪* (R^2^)coinciding with the free energy of a ruffled membrane sheet, studied previously by Leibler [49]. Away from this limit, Eq. (11) is much more involved, though, the chief reason lying in the different expansions of the square root of the metric determinant: a sheet with small ruffles of height *h* yields, in the Monge parameter-isation, 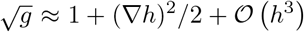 ; the ruffled tube in arc-length parametrisatio n yields 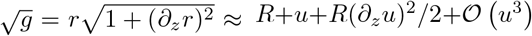 instead. The presence of terms linear in the perturbation in the arc-length parametrisation (and their absence in the Monge parameterisation) precipitates in many more terms at quadratic order in the analysis of the membrane tube.

The stability of the coated membrane tube is governed by the smallest eigenvalue 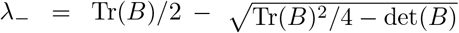 of *B* [Eq. (11)], where Tr(*B*) and det(*B*) are the trace and determinant of *B*. When *λ*_−_ *<* 0, the system is unstable to a combination of density and radius perturbations set by the elements of the corresponding eigenvector. Figure 2(a) and (b) show *λ*_−_ for 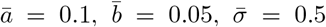, and several proteincurvature couplings 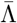 as indicated. We observe local minima at 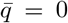 for all considered 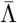 and for finite 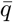 in a few cases; these minima appear, depending on 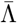, both at positive and negative *λ*_−_ values. We discern four cases: for 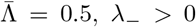, for all *q* (case I); for 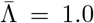, *λ*_−_ is negative only around *q* = 0 (case II); for 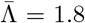 and 2.3, *λ*_−_ has a negative global minimum at *q* = 0 and a negative local minimum for *q ≠* 0 (case III); and for 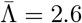, *λ*_−_ has a negative global minimum at *q ≠* 0 and a negative local minimum for *q* = 0 (case IV). In Fig. 2(c), we characterise *λ*_−_ according to the above cases in the plane of 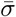 and 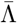. At the boundary between cases I and II, the minimum of *λ*_−_ at 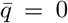 crosses zero. *λ*_−_ = 0 requires det(*B*) = 0, which, after setting 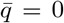 in *B*, gives the stability criterion Eq. (7) in dimensionless form, 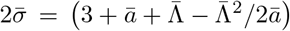. We indicate this analytical result with a red dotted line. It would be interesting also to characterise the boundary between cases II and III, where the local minimum of *λ*_−_ at 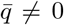 crosses zero, as this is the criterion for finite-wavelength undulations to become unstable. In this case, det(*B*) = 0 is a sixth-order polynomial in 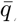 containing only even powers of 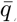, which can thus be written as a cubic polynomial, det(*B*) = *αx*^3^ + *βx*^2^ + *γx* + *δ*, with 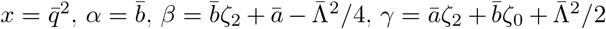, and 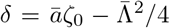. Depending on the sign of the dis-criminant Δ = *β*^2^*γ*^2^−4*αγ*^3^−4*β*^3^*δ−*27*α*^2^*δ*^2^ + 18*αβγδ*, det(*B*) = 0 thus has either 1 (if Δ *<* 0) or 3 (if Δ *>* 0) real solutions. While these expressions allow us to express the stability of the protein-covered membrane analytically, the resulting expressions are not tractable.

**Figure 2.**
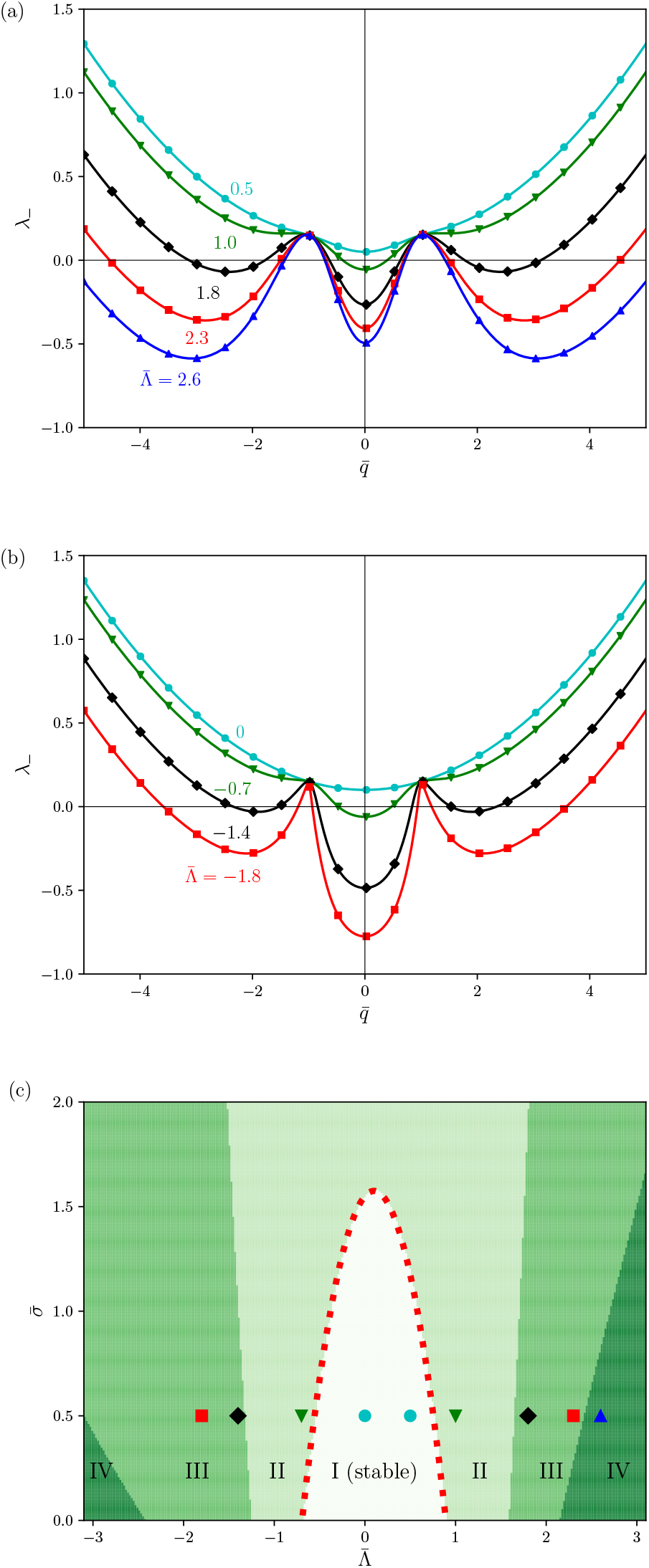
The smallest eigenvalue *λ*_−_ of *B* [Eq. (11)] for 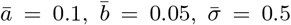, and 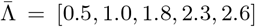 (panel a) and 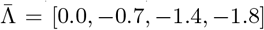 (panel b). We discern four types of minima of *λ*_−_, which we denote with I, where *λ*_−_*>* 0 for all *q*; II, where min(*λ*_−_) *<* 0 at *q* = 0 is the only minimum for which *λ*_−_ *<* 0; III, same as II, but with a secondary minimum for which *λ*_−_ *<* 0 for *q≠*0, and; IV, where min(*λ*_−_) *<* 0 at *q ≠*0. In panel (c), we characterise the minima of *λ*_−_ according to these four cases in the plane of 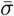 and 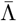. Symbols correspond to the parameter settings used in panels (a) and (b). The red dotted line represents Eq. (7).

## III. HYDRODYNAMIC INSTABILITY

Next, we consider how hydrodynamics and protein mobility set the dynamics of the membrane instability discussed above. We consider a membrane with shape operator **S** = −∇**N**, with **N** the surface normal and ∇ the surface gradient operator. The orthonormal triad{ **e**_1_, **e**_2_, **N**} provides a basis for the tangent and normal bundles on the surface. The membrane is assumed to move with a velocity **V** = **v** + *v*_*n*_**N**, to be incompressible, and to have membrane viscosity *η*_*m*_. The ambient fluid is assumed to be an incompressible Stokes fluid with viscosity *η*.

In Section S7 A of the ESI [68], we derive the following general dynamic equations. The ambient fluid obeys momentum and mass conservation equations,

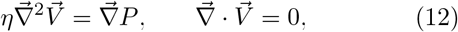

where 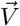 is the ambient fluid velocity and *P* is the pressure. Note that 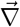 is the gradient operator in ℝ^3^.

Incompressibility on the membrane is imposed with

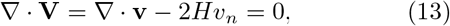

and force balance is given by

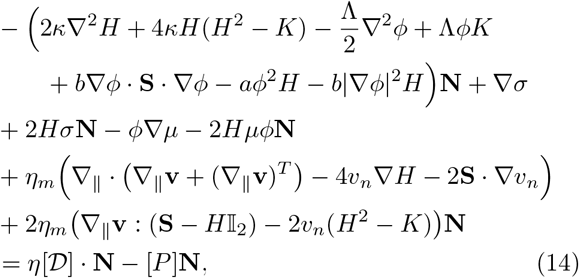

where ∇_||_**v** = ∇ **v**− (∇ **v ·N**)**N**. The tangential part of this is the covariant Stokes equation, and the normal part corresponds to the shape equation (including viscous stresses). [𝒟] is the jump in the ambient deformation rate tensor across the membrane. *K* is the Gaussian curvature.

Finally, we have a continuity equation for protein concentration given by

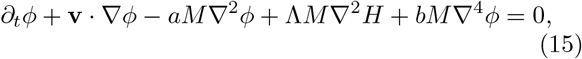

where *M* is the protein mobility on the surface (and with *∂*_*t*_ = *∂/∂t*). Equations (13)–(15) are coordinate-free versions of the equations of irreversible thermodynamics of fluid bilayers derived in Ref. [29], assuming the entropy of mixing term is expanded to quadratic order. They are closed by no-slip and no-permeation boundary conditions between the ambient fluid and membrane.

In Section S7 B of the ESI [68], we expand Eqs. (12)–(15) out to linear order for an axisymmetric tube with radius *r* = *R* + *u*(*z, t*) and protein concentration *ϕ* = Φ + *δ*Φ. Transforming to Fourier space and solving for velocities of the bulk flow and membrane, surface tension variation and pressures then gives

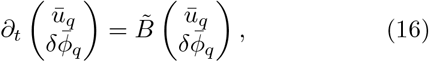

Where

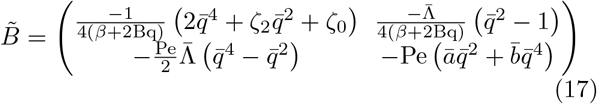

And

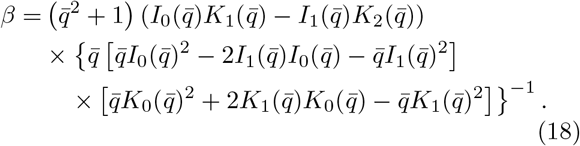

Here, Bq = *η*_*m*_*/Rη* is the Boussinesq number giving the ratio of membrane viscosity to bulk viscosity. Pe = *Mη/R*^3^Φ^2^ is the Péclet number relating surface mobility to bulk viscous advection. In the context of membranes, the Boussinesq number is sometimes called the Saffman-Delbrück number. For the case of a membrane tube, Bq*∼*1 [71], and Pe ≫ 1. Time has been non-dimensionalised by *τ* = *ηR/κ*. All other dimensionless quantities are as defined in Section II B. *I*_*m*_(*x*) and *K*_*m*_(*x*) are modified Bessel functions of the first and second kind, respectively. Note that the stability matrix 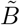 is similar to *B* [Eq. (11)] as derived from the free energy (modulo an overall minus sign), only with dissipative prefactors multiplying each row. Moreover, 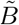 is similar to the stability matrix of a flat membrane [43] (see also Eq. (S15) of the ESI [68]), but also accounts for membrane viscosity.

To understand how hydrodynamics affects the wavelength selection of the instability, we plot the eigenvalues *λ±*of the stability matrix 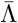 in Fig. 3(a) for Pe = Bq = 1 and several protein curvature couplings 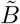. We note that, for 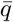, positive *λ*_*±*_ corresponds to instability, whereas the opposite is true for *B*. We see that the bulk hydrodynamics screens the small 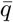 instability as the confinement of the tube gives an infinite resistance to the pearling mode in the long wavelength limit. The small 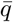 mode grows essentially as a classical pearling instability with wave-length set by the Boussinesq number [71]; see Fig. 3(b).

**Figure 3.**
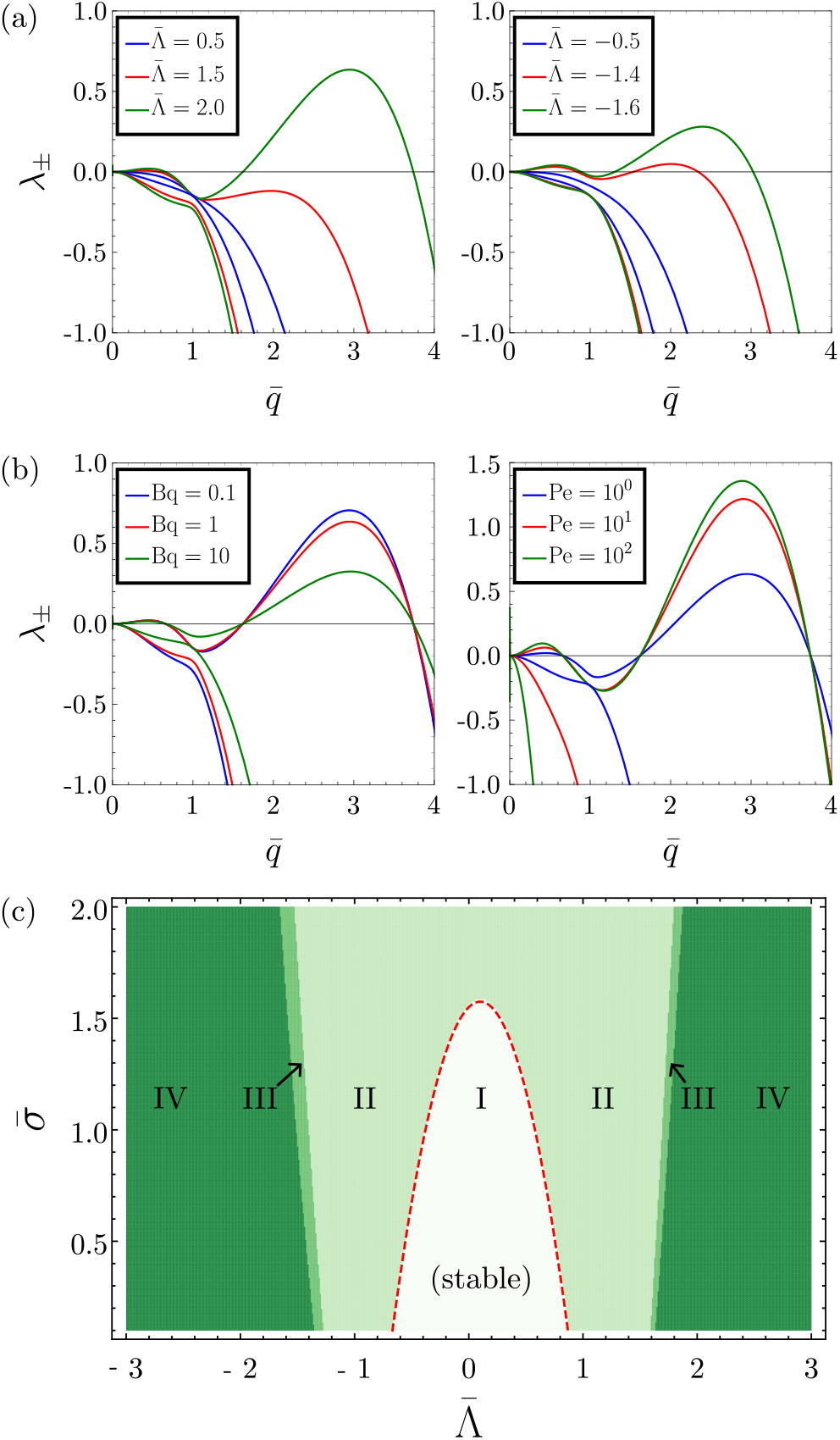
Eigenvalues *λ*_*±*_ of the dynamical stability matrix 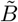 [Eq. (17)] for Bq = 1, Pe = 1, and various 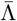 and for 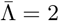 and various Bq and Pe (b). Throughout, 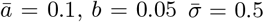. Panel (c) shows a stability diagram for Bq = 1 and Pe = 10^2^, with regions defined as in Fig. 2. The red dotted line represents Eq. (7).

The wavelength of the large 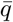 instability is relatively unchanged as the membrane dissipation controls the dissipative dynamics. Here, the length scale is essentially set by the elastic forces derived from the free energy, while the dynamical parameters control the growth rate; see Fig. 3(b). We note that for large Péclet number, the growth rate saturates to a value controlled by the membrane viscosity.

Finally, Fig. 3 shows a stability diagram similar to Fig. 2(c), now based on hydrodynamic theory. We see that the main characteristics of the instability are unchanged. Still, region III, where both high and low 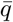 are unstable but the long wavelength instability dominates, is now heavily suppressed. Moreover, the short wavelength instability quickly becomes the fastest-growing mode, see Fig. 3(c). This analysis suggests that free energy calculations are sufficient to capture the general wavelength selection mechanisms for the dominant high 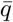 instability. Thus, we proceed with a purely energetic analysis for the paper’s final part.

## IV. ENERGY MINIMISING SHAPE

Finally, we discuss the membrane shape beyond the onset of the instability. We note that the 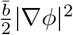 term promotes a homogeneous protein coat. The term acts as an effective line tension *γ* in the case of strong density gradi-ents between dense 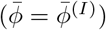 and dilute 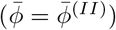 protein domains, where we estimate 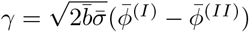 (see Sec. S8 of the ESI [68]). Therefore, we limit the discussion of tube shape to cases where the tube is covered either by a continuous protein coat or by alternating homogeneous domains of dilute or dense coats. In the latter case, the line tension *γ* acts at the interface between the dilute and dense domain. Within a homogeneous domain, the shape equations that arise from the variation of energy are solved by shapes with constant mean curvature, so-called Delaunay shapes (see ESI [68]) [72]. For all shapes discussed here, the volume is conserved. Figure 4 shows examples of the tube shapes for 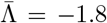 and 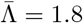. For 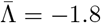, the lowest-energy shape is a weak undulation with a continuous homogeneous protein coat with a slightly higher protein density than the reference cylindrical shape. We note that there is a second solution to the shape equation characterised by an alternation of protein-free and protein-dense domains. The corresponding change in mean curvature (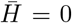 in the protein-free domain and 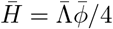 in the dense domain, with 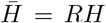) leads to a strong undulation. The energy of this tube shape is lower than that of a cylinder but higher than that of the previously discussed weak undulation with a homogeneous protein coat.

**Figure 4.**
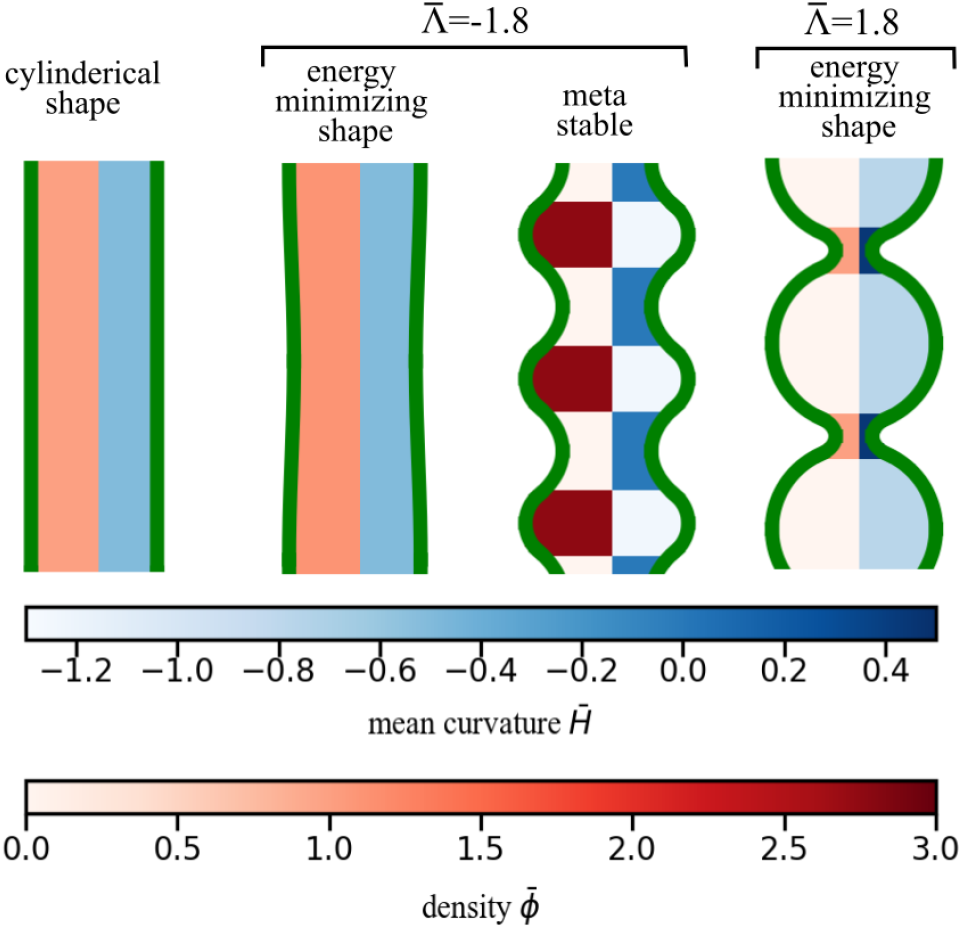
Membrane tube shape: The cylindrical shape has a constant mean curvature 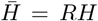, with = −1*/*2 and a constant protein density 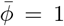. The areal protein density and mean curvature of the undulating shapes are shown as color maps. We use 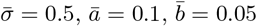, in all panels.

For 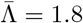, the shape of a tube can not be described by a constant and positive mean curvature. Therefore, for this case, we do not find a solution to the shape equations for a continuous protein coat. The energy-minimising shapes exhibit large undulations with extended protein-free regions. In contrast to the tube shapes for 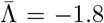, where regions with negative mean curvature exhibit a large protein density, we find the regions with negative mean curvature to be protein free.

We note the undulating shapes found in Fig. 4 and anticipated in Fig. 2 are characteristic of the small 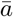 and 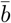 used there. For larger 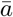 and 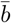, the negative local minima for 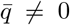 do not appear in the same 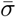 and 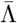 range [68]. Physically, for particles to induce a membrane tube instability, these particles need to interact strongly with the membrane, but not among themselves.

## V. CONCLUSION

We studied the stability of a membrane tube coated with proteins. By a linear stability analysis we identified short and long wavelength instabilities that we corroborated by hydrodynamic theory and through numerical calculations. Our analysis showed that both membrane and protein properties determine the onset of shape instabilities; proteins thus have a regulatory effect on the shape and stability of membrane tubes. Moreover, our model revealed an interplay between membrane shape and protein density: the membrane shape instability studied here concurred with a redistribution of proteins into a banded pattern. We hope that these results inspire experiments seeking protein bands on membrane tubes. Moreover, future work may extend our analysis to tubes with no cylindrical symmetry.

## Supporting information

List of used symbols; Fig. 2 replotted for different parameters; a discussion of instabilities of membrane sheets; derivations of Eq. 11-18.

## ACKNOWLEDGEMENTS

We thank David Andelman, Andrew Callan-Jones, Markus Deserno, and Gareth Alexander for insightful discussions. The research leading to these results has re-ceived funding from the European Union’s Horizon 2020 research and innovation programme under the Marie Sklodowska-Curie grant agreement No 801133. MJ was supported by an Advanced Grant from the European Research Council (no. 788954).

