## Supplementary material for "Stability of a biomembrane tube covered with proteins": List of used symbols; Fig. 2 replotted for different parameters; a discussion of instabilities of membrane sheets; derivations of Eq. 11-18.

### Electronic Supplementary Information to: Stability of a biomembrane tube covered with proteins

#### S1. NOTATION

Table I. Parameter, coordinates, variables

| symbol | name | (typical value/) unit | definition/source |
|---|---|---|---|
| $F_H$ | Helfrich free energy | J | Eq. (1) of the main text |
| $F_\phi$ | Ginzburg Landau free energy | J | Eq. (2) of the main text |
| $F$ | total free energy | J | $F = F_H + F_\phi$ |
| $\mathbf{S}$ | Shape Operator | $\mathrm{m}^{-1}$ | $\mathbf{S} = -\nabla \mathbf{N}$ ($\mathbf{N}$ is the surface normal) |
| $H$ | mean curvature | $\mathrm{m}^{-1}$ | $2H = \mathrm{Tr}\,(\mathbf{S})$ |
| $K$ | Gaussian curvature | $\mathrm{m}^{-2}$ | $K = \det(\mathbf{S})$ |
| $g$ | metric determinant | 1 (Monge parameterisation), $\mathrm{m}^2$ (arc length) | |
| $z$ | coordinate along tube length | m | |
| $r(z)$ | local tube radius | m | |
| $R$ | radius of a straight tube | $\sim 4 \times 10^{-6}\,\mathrm{m}$ [a] | |
| $L$ | tube length | m | |
| $\phi(z)$ | areal protein density | $\mathrm{m}^{-2}$ | |
| $\Phi$ | areal protein density | $\sim 2 \times 10^{16}\,\mathrm{m}^{-2}$ [b] | |
| $q$ | wave number | $\mathrm{m}^{-1}$ | $2\pi n/L$ |
| $u(z)$ | radius perturbation | m | $r(z) - R$ |
| $\varphi(z)$ | areal protein density perturbation | $\mathrm{m}^{-2}$ | $\phi(z) - \Phi$ |
| $\sigma$ | surface tension | $\sim 2 \times 10^{-5}\,\mathrm{J\,m}^{-2}$ | Ref. [1] |
| $\kappa$ | bending rigidity | $\sim 20k_BT = 8 \times 10^{-20}\,\mathrm{J}$ | Ref. [2] |
| $a$ | protein interactions | $\sim 7.2 \times 10^{-38}\,\mathrm{J\,m}^2$ [c] | |
| $b$ | cost of phase separating | $\mathrm{J\,m}^4$ | |
| $\Lambda$ | protein-membrane interactions | $\sim -1 \times 10^{-28}\,\mathrm{J\,m}$ [d] | |
| $\mu$ | chemical potential | J | |
| $P$ | pressure | $\mathrm{N\,m}^{-2}$ | |
| $\eta$ | shear viscosity of the fluid | $\sim 10^{-3}\,\mathrm{Pa\,s}$ | |
| $\eta_m$ | 2D membrane viscosity | $\sim 10^{-9}\,\mathrm{Pa\,m\,s}$ | Ref. [5] |
| $D$ | protein diffusion constant | $\sim 10^{-12}\,\mathrm{m}^2\,\mathrm{s}^{-1}$ | Ref. [6] |
| $M$ | protein mobility | $D/a$ | |

[a] We find this value by solving Eq. (5) of the main text for $R$ for $P = 0$, for the given values of $\Phi$ and $\kappa$, and using $a = \kappa/(R^2\Phi^2)$, see Footnote c. The stated value is slightly larger than the predicted radius $R = \sqrt{\kappa/(2\sigma)} \sim 4.5 \times 10^{-8}\,\mathrm{m}$ for an uncoated membrane tube.

[b] For this order of magnitude estimate, we considered 25% of the area of the membrane tube to be covered by proteins. The protein area of one protein is taken to be $15\,\mathrm{nm}^2$. This means we have 1 protein per $60\,\mathrm{nm}^2 \sim 1.8 \times 10^{16}\,\mathrm{m}^{-2}$.

[c] We are not aware of experimental data for this parameter. We follow [3] to come to the following estimate. Below Eq. (S13) we show that our $a$ corresponds to their $\kappa/(R^2\Phi^2) + (4T - J)/(a^2\Phi^2)$, the latter term vanishing because they set $J = 4T$. Next, we use $a = \kappa/(R^2\Phi^2)$ to solve and values for $\kappa$ and $\Phi$ to solve $R$ (see Footnote a). Inserting the found $R$ gives $a \sim 7.2 \times 10^{-38}\,\mathrm{J\,m}^2$

[d] For this order of magnitude estimate, we noted that the $-\Lambda\phi H$ term in Eq. (2) of the main text corresponds to $-4\kappa H H_0$ of Ref. [4]. Here, $H_0$ is spontaneous curvature induces by proteins. Assuming $H_0$ to be linearly dependent on the protein density, we take $H_0 \to H_0\Phi/\Phi_{\max}$, with $\Phi_{\max}$ the protein density at close packing, which we take to be around a packing fraction of 0.5, so $\Phi_{\max} \sim 3.3 \times 10^{16}\,\mathrm{m}^{-2}$; see Footnote b. We set $\phi \to \Phi$ and rewrite to $\Lambda = 4\kappa H_0/\Phi_{\max}$. Inserting $\Phi_{\max}$ and $\kappa$ and estimating the spontaneous curvature by $H_0 \sim 10^{-7}\,\mathrm{m}^{-1}$, we find $\Lambda \sim -2 \times 10^{-28}\,\mathrm{J\,m}$.

Table II. Dimensionless parameters, coordinates, and variables

| symbol | name | definition |
|---|---|---|
| $\bar{F}, \bar{F}_H, \bar{F}_\phi$ | free energies | $F/(2\pi\kappa)$ |
| $\Delta\bar{F}$ | difference free energy to free energy of a cylinder | |
| $\bar{h}$ | mean curvature | $HR$ |
| $\bar{z}$ | coordinate along tube length | $z/R$ |
| $\bar{r}$ | local tube radius | $r/R$ |
| $\bar{L}$ | tube length | $L/R$ |
| $\bar{\phi}(z)$ | areal protein density | $\phi/\Phi$ |
| $\bar{u}(\bar{z})$ | radius perturbation | $u/R$ |
| $\bar{\varphi}(\bar{z})$ | areal protein density perturbation | $\varphi/\Phi$ |
| $\bar{u}_{\bar{q}}$ | Fourier coefficient of $\bar{u}(\bar{z})$ | |
| $\bar{\varphi}_{\bar{q}}$ | Fourier coefficient of $\bar{\varphi}(\bar{z})$ | |
| $\bar{q}$ | wave number | $qR = 2\pi n/\bar{L}$ |
| $\bar{\sigma}$ | surface tension | $\sigma R^2/\kappa$ |
| $\bar{a}$ | protein interactions | $a\Phi^2 R^2/\kappa$ |
| $\bar{b}$ | cost of phase separating | $b\Phi^2/\kappa$ |
| $\bar{\Lambda}$ | protein-membrane interactions | $\Lambda\Phi R/\kappa$ |
| $\bar{\mu}$ | chemical potential | $\mu\Phi R^2/\kappa$ |
| $\bar{P}$ | pressure | $PR^3/\kappa$ |
| Bq | Boussinesq number | $\eta_m/R\eta$ |
| Pe | Péclet number | $M\eta/R^3\Phi^2$ |
| $\tau = \eta R/\kappa$ | | |

#### S2. SUPPLEMENTAL FIGURES

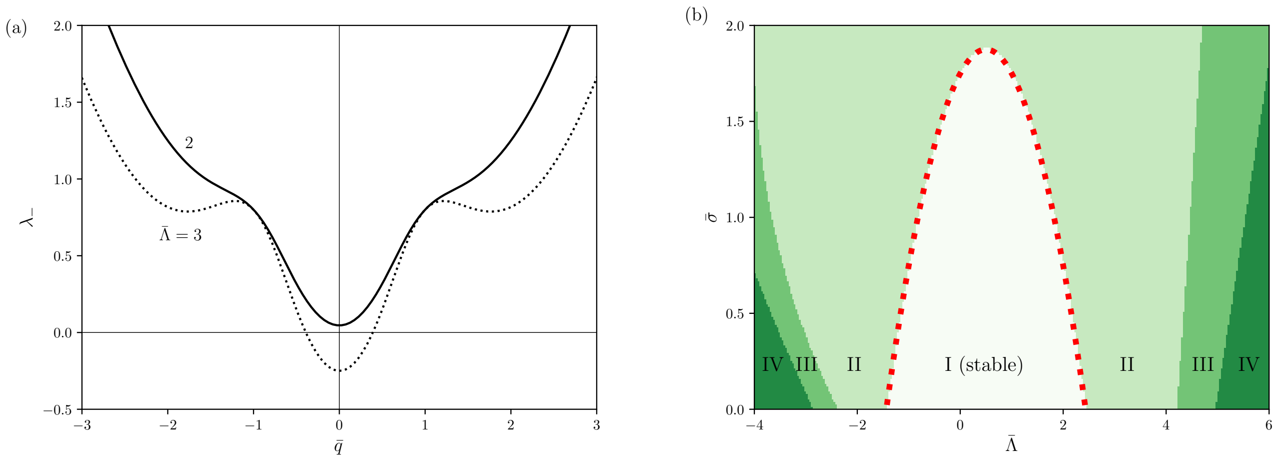


Figure S1. We redraw Fig. 2 of the main text for the larger values $\bar{a} = 0.5$ and $\bar{b} = 0.3$. Panel (a) shows the smallest eigenvalue $\lambda_-$ of $B$ [Eq. (11) of the main text] for $\bar{\sigma} = 0.5$ and $\bar{\Lambda} = 2$ and 3. Compared to Fig. 2 of the main text, the larger $\bar{a}$ and $\bar{b}$ values suppress the $q \neq 0$ minima of $\lambda_-$; hence, they suppress the generation of undulations. Panel (b) is a diagram for the stability of the membrane tube to $q = 0$ undulations. The red dotted line shows the analytical result $2\bar{\sigma} = \left(3 + \bar{a} + \bar{\Lambda} - \bar{\Lambda}^2/2\bar{a}\right)$.

#### S3. STABILITY OF A FLAT PROTEIN-COVERED MEMBRANE

We consider the same free energy $F = F_H + F_\phi$ as in Eqs. (1) and (2) of the main text and now revisit the stability of a flat protein-covered membrane sheet. In the Monge parameterisation [7], considering slight deviations $h(\mathbf{x})$ from a flat membrane, we have $\mathrm{d}A = \left(1 + \frac{1}{2}(\nabla h)^2\right)\mathrm{d}\mathbf{x}$ and $H = \frac{1}{2}\nabla^2 h$, where $\nabla$ is the gradient operator. We can then write $F = F_H + F_\phi$ with

$$F = \frac{1}{2}\int \mathrm{d}\mathbf{x}\, \left(1 + \frac{1}{2}(\nabla h)^2\right) \left\{2\sigma + \kappa(\nabla^2 h)^2 - \Lambda\phi(\nabla^2 h) - 2\mu\phi + a\phi^2 + b(\nabla\phi)^2\right\}\,. \tag{S1}$$

Inserting

$$h(\mathbf{x}) = \sum_{\mathbf{q}\neq\mathbf{0}} h_{\mathbf{q}} \mathrm{e}^{\mathrm{i}\mathbf{q}\cdot\mathbf{x}}, \qquad \phi(\mathbf{x}) = \Phi + \sum_{\mathbf{q}\neq\mathbf{0}} \phi_{\mathbf{q}} \mathrm{e}^{\mathrm{i}\mathbf{q}\cdot\mathbf{x}} \equiv \Phi + \varphi(\mathbf{x})\,, \tag{S2}$$

into Eq. (S1) yields

$$\begin{aligned} F = \frac{1}{2}\int \mathrm{d}\mathbf{x}\, &\left(1 - \frac{1}{2}\sum_{\mathbf{q},\mathbf{q}'\neq\mathbf{0}} \mathbf{q}\cdot\mathbf{q}' h_{\mathbf{q}} h_{\mathbf{q}'} \mathrm{e}^{\mathrm{i}(\mathbf{q}+\mathbf{q}')\cdot\mathbf{x}}\right) \times \\ &\times \Bigg\{2\sigma + \kappa \sum_{\mathbf{q},\mathbf{q}'\neq\mathbf{0}} q^2 (q')^2 h_{\mathbf{q}} h_{\mathbf{q}'} \mathrm{e}^{\mathrm{i}(\mathbf{q}+\mathbf{q}')\cdot\mathbf{x}} - \Lambda \left(\Phi + \sum_{\mathbf{q}\neq\mathbf{0}} \phi_{\mathbf{q}} \mathrm{e}^{\mathrm{i}\mathbf{q}\cdot\mathbf{x}}\right) \left(-\sum_{\mathbf{q}'\neq\mathbf{0}} (q')^2 h_{\mathbf{q}'} \mathrm{e}^{\mathrm{i}\mathbf{q}'\cdot\mathbf{x}}\right) - 2\mu \left(\Phi + \sum_{\mathbf{q}\neq\mathbf{0}} \phi_{\mathbf{q}} \mathrm{e}^{\mathrm{i}\mathbf{q}\cdot\mathbf{x}}\right) \\ &\quad + a \left(\Phi^2 + 2\sum_{\mathbf{q}\neq\mathbf{0}} \phi_{\mathbf{q}} \mathrm{e}^{\mathrm{i}\mathbf{q}\cdot\mathbf{x}} + \sum_{\mathbf{q},\mathbf{q}'\neq\mathbf{0}} \phi_{\mathbf{q}}\phi_{\mathbf{q}'} \mathrm{e}^{\mathrm{i}(\mathbf{q}+\mathbf{q}')\cdot\mathbf{x}}\right) - b \left(\sum_{\mathbf{q},\mathbf{q}'\neq\mathbf{0}} \mathbf{q}\cdot\mathbf{q}' \phi_{\mathbf{q}}\phi_{\mathbf{q}'} \mathrm{e}^{\mathrm{i}(\mathbf{q}+\mathbf{q}')\cdot\mathbf{x}}\right)^2 \Bigg\}\,, \end{aligned} \tag{S3}$$

with $q = |\mathbf{q}|$. We can partition the above expression as $F = F_0 + F_1 + F_2$, with $F_0 = \mathcal{O}\left(\epsilon^0\right)$, $F_1 = \mathcal{O}\left(\epsilon^1\right)$, $F_2 = \mathcal{O}\left(\epsilon^2\right)$ and $\epsilon$ shorthand for the perturbations $h_{\mathbf{q}}$ and $\phi_{\mathbf{q}}$. The $F_0$ term provides a constant offset to the free energy and can be ignored from hereon. The $F_1$ term does not contribute as it is of the form

$$F_1 = \int \mathrm{d}\mathbf{x} \sum_{\mathbf{q}\neq\mathbf{0}} \mathrm{e}^{\mathrm{i}\mathbf{q}\cdot\mathbf{x}} \left[h_{\mathbf{q}}(\ldots) + \phi_{\mathbf{q}}(\ldots)\right] = \sum_{\mathbf{q}\neq\mathbf{0}} \delta_{\mathbf{q},0} \left[h_{\mathbf{q}}(\ldots) + \phi_{\mathbf{q}}(\ldots)\right] = 0\,. \tag{S4}$$

We thus find that $F$ is given, up to second order in the perturbations $h_{\mathbf{q}}$ and $\phi_{\mathbf{q}}$, by

$$F = \frac{1}{2}\int \mathrm{d}\mathbf{x} \sum_{\mathbf{q},\mathbf{q}'\neq\mathbf{0}} \mathrm{e}^{\mathrm{i}(\mathbf{q}+\mathbf{q}')\cdot\mathbf{x}} \left\{\left[\left(\mu\Phi - \sigma - \frac{a}{2}\Phi^2\right)\mathbf{q}\cdot\mathbf{q}' + \kappa q^2 (q')^2\right] h_{\mathbf{q}} h_{\mathbf{q}'} + \Lambda (q')^2 \phi_{\mathbf{q}} h_{\mathbf{q}'} + (a - b\mathbf{q}\cdot\mathbf{q}')\phi_{\mathbf{q}}\phi_{\mathbf{q}'}\right\}\,. \tag{S5}$$

Integrating over $\mathbf{x}$ gives

$$\begin{aligned} F &= \frac{A}{2} \sum_{\mathbf{q},\mathbf{q}'\neq\mathbf{0}} \delta_{\mathbf{q},-\mathbf{q}'} \left\{\left[\left(\mu\Phi - \sigma - \frac{a}{2}\Phi^2\right)\mathbf{q}\cdot\mathbf{q}' + \kappa q^2 (q')^2\right] h_{\mathbf{q}} h_{\mathbf{q}'} + \Lambda (q')^2 \phi_{\mathbf{q}} h_{\mathbf{q}'} + (a - b\mathbf{q}\cdot\mathbf{q}')\phi_{\mathbf{q}}\phi_{\mathbf{q}'}\right\} \\ &= \frac{A}{2} \sum_{\mathbf{q}\neq\mathbf{0}} \left\{\left[\left(\sigma + \frac{a}{2}\Phi^2 - \mu\Phi\right) q^2 + \kappa q^4\right] h_{\mathbf{q}} h_{-\mathbf{q}} + \Lambda q^2 \phi_{\mathbf{q}} h_{-\mathbf{q}} + (a + bq^2)\phi_{\mathbf{q}}\phi_{-\mathbf{q}}\right\}\,, \end{aligned} \tag{S6}$$

with $A$ the area of the membrane patch. With the Euler Lagrange equation $\delta F/\delta\phi = \nabla\cdot(\delta F/\delta\nabla\phi)$ we find $-\Lambda(\nabla^2 h) - 2\mu + 2a\phi = 2b\nabla^2\phi$. For a homogeneously-coated flat membrane, this yields $\mu = a\Phi$. We can thus write Eq. (S6) in matrix form as

$$F = \frac{A}{2} \sum_{\mathbf{q}\neq\mathbf{0}} \begin{pmatrix} h_{\mathbf{q}} & \phi_{\mathbf{q}} \end{pmatrix} \begin{pmatrix} \kappa q^4 + \left(\sigma - \frac{a}{2}\Phi^2\right) q^2 & \frac{1}{2}\Lambda q^2 \\ \frac{1}{2}\Lambda q^2 & a + bq^2 \end{pmatrix} \begin{pmatrix} h_{-\mathbf{q}} \\ \phi_{-\mathbf{q}} \end{pmatrix}\,. \tag{S7}$$

For small $q$, the eigenvalues of the above matrix read

$$\lambda_- = \left(\sigma - \frac{a}{2}\Phi^2\right) q^2 + \left(\kappa - \frac{\Lambda^2}{4a}\right) q^4 + \mathcal{O}\left(q^5\right)\,, \qquad \lambda_+ = a + bq^2 + \frac{\Lambda^2}{4a} q^4 + \mathcal{O}\left(q^5\right)\,, \tag{S8}$$

from which we see that the flat membrane becomes unstable when $\sigma < a\Phi^2/2$. For the case $\sigma = a\Phi^2/2$, we recover the instability criterion $\kappa < \Lambda^2/(4a)$ of Ref. [8] (there is a factor 2 difference in our definitions of $\Lambda$, as Ref. [8] defines the mean curvature in terms of principle curvatures ($R_1$ and $R_2$) as $H = 1/R_1 + 1/R_2$, without a factor 2.)

Equation (S8) implies the equilibrium height variance $\langle|h_{\mathbf{q}}|^2\rangle = k_B T/\lambda_-$, see Eqs. (8) and (9) of Ref. [9]. In other words, the long-wavelength height fluctuations diverge at the stated instability criterion.

In the above derivation, we followed Refs. [10–12], who included the $\sqrt{g} \sim 1+\frac{1}{2}(\nabla h)^2$ prefactor for the $\phi$-dependent terms of $F$. In contrast, Refs. [8, 9, 13–15] multiplied the area element $\sqrt{g}$ only with the term proportional to $\sigma$, in which case the free energy reads

$$F = \frac{1}{2}\int \mathrm{d}\mathbf{x}\,\left\{\sigma(\nabla h)^2 + \kappa(\nabla^2 h)^2 - \Lambda\phi(\nabla^2 h) - 2\mu\phi + a\phi^2 + b(\nabla\phi)^2\right\}\,, \tag{S9}$$

instead of Eq. (S1). Repeating the above derivation, we now find

$$F = \frac{A}{2}\sum_{\mathbf{q}\neq\mathbf{0}} \begin{pmatrix} h_{\mathbf{q}} & \phi_{\mathbf{q}} \end{pmatrix} \begin{pmatrix} \kappa q^4 + \sigma q^2 & \frac{1}{2}\Lambda q^2 \\ \frac{1}{2}\Lambda q^2 & a + bq^2 \end{pmatrix} \begin{pmatrix} h_{-\mathbf{q}} \\ \phi_{-\mathbf{q}} \end{pmatrix}\,, \tag{S10}$$

and eigenvalues

$$\lambda_- = \sigma q^2 + \left(\kappa - \frac{\Lambda^2}{4a}\right) q^4 + \mathcal{O}\left(q^5\right)\,, \qquad \lambda_+ = a + bq^2 + \frac{\Lambda^2}{4a} q^4 + \mathcal{O}\left(q^5\right)\,. \tag{S11}$$

In this case, the flat membrane becomes unstable when $\sigma < 0$.

Finally, we compare the above expressions to the free energy of Veksler and Gov [3],

$$F = \int \mathrm{d}\mathbf{x}\,\left\{\frac{1}{2}(\sigma - \alpha\bar{\phi})(\nabla h)^2 + \frac{\kappa}{2}\left(\nabla^2 h + \frac{\bar{\phi}}{R}\right)^2 + \frac{T}{a^2}[\bar{\phi}\ln\bar{\phi} + (1-\bar{\phi})\ln(1-\bar{\phi})] + \frac{J}{2a^2}\bar{\phi}(1-\bar{\phi}) + \frac{J}{4}(\nabla\bar{\phi})^2\right\}\,, \tag{S12}$$

whose various parameters we will relate to the parameters in Eq. (S1). Reference [3] considers Eq. (S12) around $\bar{\phi} = 1/2$, so we substitute $\bar{\phi} \to 1/2 + \bar{\phi}$ and expand for small $\bar{\phi} \ll 1$, giving

$$\begin{aligned} F = &\int \mathrm{d}\mathbf{x}\left\{\frac{1}{2}\left(\sigma - \frac{\alpha}{2} - \alpha\bar{\phi}\right)(\nabla h)^2 + \frac{\kappa}{2}\left[(\nabla^2 h)^2 + \frac{1+2\bar{\phi}}{R}\nabla^2 h + \frac{1}{4R} + \frac{\bar{\phi}}{R^2} + \frac{\bar{\phi}^2}{R^2}\right]\right. \\ &\left. + \frac{T}{a^2}\left(-\ln 2 + 2\bar{\phi}^2 + \frac{4}{3}\bar{\phi}^4\right) + \frac{J}{2a^2}\left(\frac{1}{4} - \bar{\phi}^2\right) + \frac{J}{4}(\nabla\bar{\phi})^2\right\} \\ = &\, F_0 + \frac{1}{2}\int \mathrm{d}\mathbf{x}\left\{\left(\sigma - \frac{\alpha}{2}\right)(\nabla h)^2 + \kappa\left(\nabla^2 h\right)^2 + \frac{2\kappa}{R}\bar{\phi}\nabla^2 h + \frac{\kappa}{R^2}\bar{\phi} + \left(\frac{\kappa}{R^2} + \frac{4T-J}{a^2}\right)\bar{\phi}^2 + \frac{J}{2}(\nabla\bar{\phi})^2\right. \\ &\left. - \alpha\bar{\phi}(\nabla h)^2 + \frac{\kappa}{R}\nabla^2 h + \frac{2}{3}\frac{T}{a^2}\bar{\phi}^4\right\}, \end{aligned} \tag{S13}$$

where $F_0$ is a constant offset to the free energy that we ignore from hereon. We see that the first line of Eq. (S13) coincides with Eq. (S9) if we take $\sigma \to \sigma - \alpha/2$, $\Lambda \to -2\kappa/(R\Phi)$, $\mu \to -\kappa/(2R^2\Phi)$, $a \to \kappa/(R^2\Phi^2) + (4T-J)/(a^2\Phi^2)$ and $b \to J/(2\Phi^2)$ in the latter expression. The second line of Eq. (S13) contains terms not in Eq. (S9). We can again substitute the Fourier represented perturbed height and protein density Eq. (S2), where we should set $\Phi = 0$ as we already assumed that $\bar{\phi} \ll 1$. This yields

$$F = \frac{1}{2}\int \mathrm{d}\mathbf{x} \sum_{\mathbf{q},\mathbf{q}'\neq\mathbf{0}} \mathrm{e}^{\mathrm{i}(\mathbf{q}+\mathbf{q}')\cdot\mathbf{x}} \left\{\left[\left(\frac{\alpha}{2} - \sigma\right)\mathbf{q}\cdot\mathbf{q}' + \kappa q^2 (q')^2\right] h_{\mathbf{q}} h_{\mathbf{q}'} - \frac{2\kappa}{R}(q')^2 \bar{\phi}_{\mathbf{q}} h_{\mathbf{q}'} + \left(\frac{\kappa}{R} + \frac{4T-J}{a^2} - \frac{J}{2}\mathbf{q}\cdot\mathbf{q}'\right)\bar{\phi}_{\mathbf{q}}\bar{\phi}_{\mathbf{q}'}\right\}, \tag{S14}$$

where, notably, the second line of Eq. (S13) does not contribute as it does not yield terms of second order in the perturbations. In matrix form, we find

$$F = \frac{A}{2}\sum_{\mathbf{q}\neq\mathbf{0}} \begin{pmatrix} h_{\mathbf{q}} & \bar{\phi}_{\mathbf{q}} \end{pmatrix} \begin{pmatrix} \kappa q^4 + \left(\sigma - \frac{\alpha}{2}\right) q^2 & -\frac{\kappa}{R} q^2 \\ -\frac{\kappa}{R} q^2 & \frac{\kappa}{R} + \frac{4T-J}{a^2} + \frac{J}{2} q^2 \end{pmatrix} \begin{pmatrix} h_{-\mathbf{q}} \\ \bar{\phi}_{-\mathbf{q}} \end{pmatrix}\,. \tag{S15}$$

The matrix in Eq. (S15) is similar but not equal to the matrix $L$ in Eq. (8) of Ref. [3], which they studied to determine the flat membrane's stability. We ignore the protrusion force $f$ of Ref. [3], and set $\bar{\phi}_0 = 0$ in their $L$. Then, comparing their $L$ to the matrix in Eq. (S15), the most important difference is an overall factor $D\eta q^2/T$ that multiplies the lower row of the matrix $L$. This factor stems from $L$ being derived from a hydrodynamic equation containing the viscosity $\eta$ and a protein continuity equation containing the diffusion constant of proteins $D$; the continuity equation has two more nabla operators than the hydrodynamic equation, which explains the factor $q^2$. Importantly, this overall factor difference between the two rows of $L$ and the matrix in Eq. (S15) means that these matrices have different eigenvalues and yield different instability criteria.

#### S4. STABILITY OF AN UNCOATED CYLINDRICAL VESICLE

Here, we rederive results for the stability of an uncoated cylindrical vesicle. We split the dimensionless free energy [Eq. (9) of the main text] into $\bar{F}_H = \bar{F}_g + \bar{F}_{\bar{\sigma}} + \bar{F}_{\bar{P}}$ with

$$\bar{F}_g = \frac{1}{2}\int_0^{\bar{L}} \mathrm{d}\bar{z}\, \frac{\bar{r}}{\sqrt{1+(\partial_{\bar{z}}\bar{r})^2}} \left[\frac{\partial_{\bar{z}}^2 \bar{r}}{1+(\partial_{\bar{z}}\bar{r})^2} - \frac{1}{\bar{r}}\right]^2 , \tag{S16a}$$

$$\bar{F}_{\bar{\sigma}} = \bar{\sigma}\int_0^{\bar{L}} \mathrm{d}\bar{z}\, \bar{r}\sqrt{1+(\partial_{\bar{z}}\bar{r})^2}\,, \tag{S16b}$$

$$\bar{F}_{\bar{P}} = \frac{1}{2}\bar{P}\int_0^{\bar{L}} \mathrm{d}\bar{z}\, \bar{r}^2 . \tag{S16c}$$

We want to know whether a cylinder ($\bar{r}=1$) is stable to small perturbations ($\bar{u} \ll 1$) of its radius $\bar{r}(\bar{z}) = 1 + \bar{u}(\bar{z})$. Inserting $\bar{r}(\bar{z}) = 1+\bar{u}(\bar{z})$ into Eq. (S16a) (with $\bar{u}(\bar{z})$ given in Eq. (10) of the main text) yields

$$\bar{F}_g = \frac{1}{2}\int_0^{\bar{L}} \mathrm{d}\bar{z}\, \frac{1+\bar{u}}{\sqrt{1+(\partial_{\bar{z}}\bar{u})^2}} \left[\frac{\partial_{\bar{z}}^2 \bar{u}}{1+(\partial_{\bar{z}}\bar{u})^2} - \frac{1}{1+\bar{u}}\right]^2 . \tag{S17}$$

Terms linear $\bar{u}$ vanish for the same reason as discussed for Eq. (S4). We thus focus on terms up to $\mathcal{O}(\bar{u}^2)$. At that order, $\bar{F}_g$ reads

$$\bar{F}_g = \frac{1}{2}\int_0^{\bar{L}} \mathrm{d}\bar{z}\,(1+\bar{u})\left(1-\frac{(\partial_{\bar{z}}\bar{u})^2}{2}\right)\left\{\partial_{\bar{z}}^2\bar{u}\left[1-\mathcal{O}(\bar{u}^2)\right] - \left[1-\bar{u}+\bar{u}^2+\mathcal{O}(\bar{u}^3)\right]\right\}^2 . \tag{S18}$$

Writing out the integrand and focussing on $\mathcal{O}(\bar{u}^2)$ terms,

$$\begin{aligned}
&\left[(1+\bar{u})\left(1-\frac{(\partial_{\bar{z}}\bar{u})^2}{2}\right)\right]\left(\partial_{\bar{z}}^2\bar{u}\left[1-\mathcal{O}(\bar{u}^2)\right] - \left[1-\bar{u}+\bar{u}^2+\mathcal{O}(\bar{u}^3)\right]\right)^2 \\
&= \left[1+\bar{u}-\frac{(\partial_{\bar{z}}\bar{u})^2}{2}+\mathcal{O}(\bar{u}^3)\right]\left(1-2(\bar{u}+\partial_{\bar{z}}^2\bar{u})+2\bar{u}^2+(\bar{u}+\partial_{\bar{z}}^2\bar{u})^2+\mathcal{O}(\bar{u}^3)\right) \\
&= \ldots + 2\bar{u}^2 + (\bar{u}+\partial_{\bar{z}}^2\bar{u})^2 - 2\left(\bar{u}^2+\bar{u}\partial_{\bar{z}}^2\bar{u}\right) - \frac{(\partial_{\bar{z}}\bar{u})^2}{2} + \mathcal{O}(\bar{u}^3) \\
&= \ldots + \sum_{\bar{q},\bar{q}'\neq 0} \bar{u}_{\bar{q}}\bar{u}_{\bar{q}'}\mathrm{e}^{\mathrm{i}(\bar{q}+\bar{q}')\bar{z}}\left\{2+1-2\bar{q}^2+\bar{q}^2(\bar{q}')^2-2\left(1-\bar{q}^2\right)+\frac{\bar{q}\bar{q}'}{2}\right\} + \mathcal{O}(\bar{u}^3)\,,
\end{aligned} \tag{S19}$$

where we used $\partial_{\bar{z}}\bar{r} = \partial_{\bar{z}}\bar{u} = \sum_{\bar{q}\neq 0}\mathrm{i}\bar{q}\bar{u}_{\bar{q}}\mathrm{e}^{\mathrm{i}\bar{q}\bar{z}}$ and $\partial_{\bar{z}}^2\bar{r} = \partial_{\bar{z}}^2\bar{u} = -\sum_{\bar{q}\neq 0}\bar{q}\bar{u}_{\bar{q}}\mathrm{e}^{\mathrm{i}\bar{q}\bar{z}}$ in the last step. The dots represent terms linear in $\bar{u}$. We thus find

$$\bar{F}_g = \frac{1}{2}\int_0^{\bar{L}} \mathrm{d}\bar{z} \sum_{\bar{q},\bar{q}'\neq 0} \bar{u}_{\bar{q}}\bar{u}_{\bar{q}'}\mathrm{e}^{\mathrm{i}(\bar{q}+\bar{q}')\bar{z}}\left(1+\bar{q}^2(\bar{q}')^2+\frac{\bar{q}\bar{q}'}{2}\right) = \frac{\bar{L}}{2}\sum_{\bar{q}\neq 0}\bar{u}_{\bar{q}}\bar{u}_{-\bar{q}}\left(\bar{q}^4-\frac{1}{2}\bar{q}^2+1\right). \tag{S20}$$

Next, inserting $\bar{r}(\bar{z}) = 1+\bar{u}(\bar{z})$ into $\bar{F}_{\bar{P}}$ [Eq. (S16c)] gives, at $\mathcal{O}(\bar{u}^2)$,

$$\bar{F}_{\bar{P}} = -\frac{\bar{P}}{2}\int_0^{\bar{L}} \mathrm{d}\bar{z}\left(1+\sum_{\bar{q}\neq 0}\bar{u}_{\bar{q}}\mathrm{e}^{\mathrm{i}\bar{q}\bar{z}}\right)^2 = \ldots - \frac{\bar{P}}{2}\int_0^{\bar{L}} \mathrm{d}\bar{z}\sum_{\bar{q},\bar{q}'\neq 0}\bar{u}_{\bar{q}}\bar{u}_{\bar{q}'}\mathrm{e}^{\mathrm{i}(\bar{q}+\bar{q}')\bar{z}} = -\frac{\bar{P}\bar{L}}{2}\sum_{\bar{q}\neq 0}\bar{u}_{\bar{q}}\bar{u}_{-\bar{q}}\,, \tag{S21}$$

where the lower dots in the second line refer to $\mathcal{O}(\bar{u}^0)$ and $\mathcal{O}(\bar{u}^1)$ terms.

Finally, the surface tension term Eq. (S16b) reads, at $\mathcal{O}(\bar{u}^2)$,

$$\begin{aligned}
\bar{F}_{\bar{\sigma}} &= \bar{\sigma}\int_0^{\bar{L}} \mathrm{d}\bar{z}\,(1+\bar{u})\left(1+\frac{(\partial_{\bar{z}}\bar{u})^2}{2}+\mathcal{O}(\bar{u}^4)\right) = \cdots + \bar{\sigma}\int_0^{\bar{L}} \mathrm{d}\bar{z}\,\frac{(\partial_{\bar{z}}\bar{u})^2}{2} + \mathcal{O}(\bar{u}^3) \\
&= \frac{\bar{\sigma}}{2}\int_0^{\bar{L}} \mathrm{d}\bar{z}\sum_{\bar{q},\bar{q}'\neq 0}(\mathrm{i})^2\bar{q}\bar{q}'\bar{u}_{\bar{q}}\bar{u}_{\bar{q}'}\mathrm{e}^{\mathrm{i}(\bar{q}+\bar{q}')\bar{z}} \\
&= \frac{\bar{\sigma}\bar{L}}{2}\sum_{\bar{q}\neq 0}\bar{q}^2\bar{u}_{\bar{q}}\bar{u}_{-\bar{q}}\,.
\end{aligned} \tag{S22}$$

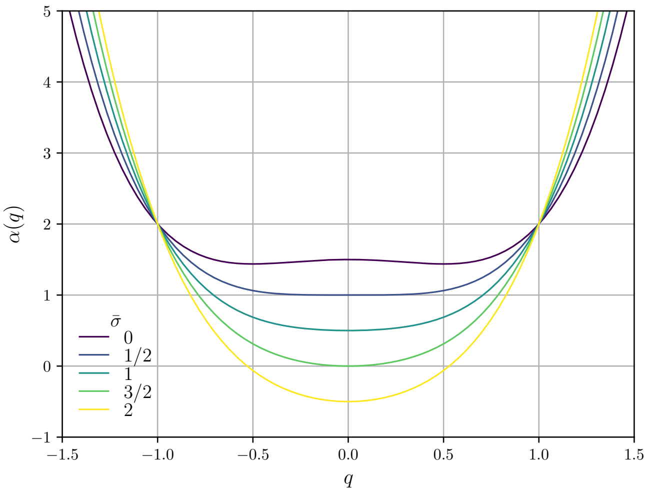


Figure S2. Plot of $\alpha(\bar{q})$ [Eq. (S24)]. The cylindrical vesicle becomes unstable ($\alpha < 0$) for perturbations at a wavelength $\bar{q} = 0$ for $\bar{\sigma} > 3/2$.

Collecting Eqs. (S20)–(S22) now yields

$$\bar{F}_H = \frac{\bar{L}}{4} \sum_{\bar{q} \neq 0} \bar{u}_{\bar{q}} \bar{u}_{-\bar{q}} \left\{ 2\bar{q}^4 + \bar{q}^2 \left( 2\bar{\sigma} - 1 \right) + 2 - 2\bar{P} \right\} . \tag{S23}$$

Inserting the Laplace pressure $\bar{P} = \bar{\sigma} - 1/2$ [the first terms of Eq. (5) of the main text] into Eq. (S23) gives [16–19]

$$\bar{F}_H = \frac{\bar{L}}{2} \sum_{\bar{q} \neq 0} \bar{u}_{\bar{q}} \bar{u}_{-\bar{q}} \underbrace{\bar{\sigma} \left( \bar{q}^2 - 1 \right) + \bar{q}^4 - \frac{1}{2}\bar{q}^2 + \frac{3}{2}}_{\alpha(\bar{q})} . \tag{S24}$$

We find that $\alpha(\bar{q})$ has minima at

$$\partial_{\bar{q}} \alpha(\bar{q}) = \bar{q} \left( 4\bar{q}^2 + 2\bar{\sigma} - 1 \right) = 0 \qquad \Rightarrow \qquad \bar{q} = 0 \wedge \bar{q} = \pm \frac{1}{2} \sqrt{1 - 2\bar{\sigma}} \, , \tag{S25}$$

see also Fig. S2, where we plot $\alpha(q)$ as defined in Eq. (S24). The cylinder is stable if $\alpha(q = 0) = -\bar{\sigma} + 3/2 > 0$; hence $\bar{\sigma} < 3/2$. In terms of the original variables, this reads $\sigma R^2/\kappa < 3/2$; hence $R < \sqrt{3\kappa/(2\sigma)}$.

#### S5. STABILITY OF AN COATED CYLINDRICAL VESICLE [DERIVATION OF EQ. (11) OF THE MAIN TEXT]

Of the free energy $\bar{F} = \bar{F}_H + \bar{F}_\phi$ [Eq. (9) of the main text] of a cylindrical vesicle coated with proteins, we considered the membrane instabilities associated with $\bar{F}_H$ in Section S4. We partition the other term into $\bar{F}_\phi = \bar{F}_{\bar{\Lambda}} + \bar{F}_{\bar{\mu}} + \bar{F}_{\bar{a}} + \bar{F}_{\bar{b}}$ where

$$\bar{F}_{\bar{\Lambda}} = -\frac{\bar{\Lambda}}{2} \int_0^{\bar{L}} \mathrm{d}\bar{z} \, \bar{\phi} \left( \frac{\bar{r} \partial_{\bar{z}}^2 \bar{r}}{1 + (\partial_{\bar{z}} \bar{r})^2} - 1 \right) , \tag{S26a}$$

$$\bar{F}_{\bar{\mu}} = -\bar{\mu} \int_0^{\bar{L}} \mathrm{d}\bar{z} \, \bar{r} \sqrt{1 + (\partial_{\bar{z}} \bar{r})^2} \bar{\phi}, \tag{S26b}$$

$$\bar{F}_{\bar{a}} = \frac{\bar{a}}{2} \int_0^{\bar{L}} \mathrm{d}\bar{z} \, \bar{r} \sqrt{1 + (\partial_{\bar{z}} \bar{r})^2} \bar{\phi}^2, \tag{S26c}$$

$$\bar{F}_{\bar{b}} = \frac{\bar{b}}{2} \int_0^{\bar{L}} \mathrm{d}\bar{z} \, \bar{r} \sqrt{1 + (\partial_{\bar{z}} \bar{r})^2} (\partial_{\bar{z}} \bar{\phi})^2. \tag{S26d}$$

We insert $\bar{r}(\bar{z}) = 1 + \bar{u}(\bar{z})$ and $\bar{\phi} = 1 + \bar{\varphi}(\bar{z})$ (with $\bar{u}(\bar{z})$ and $\bar{\varphi}(\bar{z})$ given in Eq. (10) of the main text) in each of these terms. Again, constant terms cannot affect any physical observable and terms linear in the perturbations drop for the reason shown in Eq. (S4). Focussing on the quadratic terms, we find

$$\begin{aligned}
\bar{F}_{\bar{\Lambda}} &= -\frac{\bar{\Lambda}}{2}\int_0^{\bar{L}} \mathrm{d}\bar{z}\ (1+\bar{\varphi})\left\{(1+\bar{u})\,\partial_{\bar{z}}^2\bar{u}\left[1+\mathcal{O}\left(\bar{u}^2\right)\right]-1\right\} \\
&= \mathcal{O}\left(\bar{u}^0,\bar{u}^1,\bar{\varphi}^0,\bar{\varphi}^1\right) - \frac{\bar{\Lambda}}{2}\int_0^{\bar{L}} \mathrm{d}\bar{z}\ \left(\bar{\varphi}+\bar{u}\right)\partial_{\bar{z}}^2\bar{u} \\
&= \frac{\bar{\Lambda}}{2}\int_0^{\bar{L}} \mathrm{d}\bar{z} \sum_{\bar{q},\bar{q}'\neq 0}\left(\bar{\phi}_{\bar{q}}+\bar{u}_{\bar{q}}\right)(\bar{q}')^2\bar{u}_{\bar{q}'}\mathrm{e}^{\mathrm{i}(\bar{q}+\bar{q}')\bar{z}} \\
&= \frac{\bar{L}\bar{\Lambda}}{2}\sum_{\bar{q}\neq 0}\bar{q}^2\left(\bar{u}_{\bar{q}}\bar{u}_{-\bar{q}}+\bar{\phi}_{\bar{q}}\bar{u}_{-\bar{q}}\right)\,,
\end{aligned} \tag{S27}$$

$$\begin{aligned}
\bar{F}_{\bar{\mu}} &= -\bar{\mu}\int_0^{\bar{L}} \mathrm{d}\bar{z}\ (1+\bar{u})\left(1+\frac{(\partial_{\bar{z}}\bar{u})^2}{2}\right)(1+\bar{\varphi}) = -\bar{\mu}\int_0^{\bar{L}} \mathrm{d}\bar{z}\ \left(\bar{u}\bar{\varphi}+\frac{(\partial_{\bar{z}}\bar{u})^2}{2}\right) \\
&= -\bar{\mu}\int_0^{\bar{L}} \mathrm{d}\bar{z} \sum_{\bar{q},\bar{q}'\neq 0}\mathrm{e}^{\mathrm{i}(\bar{q}+\bar{q}')\bar{z}}\left(\bar{\phi}_{\bar{q}}\bar{u}_{\bar{q}'}-\bar{u}_{\bar{q}}\bar{u}_{\bar{q}'}\frac{\bar{q}\bar{q}'}{2}\right) \\
&= -\frac{\bar{L}\bar{\mu}}{2}\sum_{\bar{q}\neq 0}2\bar{\phi}_{\bar{q}}\bar{u}_{-\bar{q}}+\bar{u}_{\bar{q}}\bar{u}_{-\bar{q}}\bar{q}^2\,,
\end{aligned} \tag{S28}$$

$$\begin{aligned}
\bar{F}_{\bar{a}} &= \frac{\bar{a}}{2}\int_0^{\bar{L}} \mathrm{d}\bar{z}\ (1+\bar{u})\left(1+\frac{(\partial_{\bar{z}}\bar{u})^2}{2}\right)\left(1+2\bar{\varphi}+\bar{\varphi}^2\right) \\
&= \frac{\bar{a}}{2}\int_0^{\bar{L}} \mathrm{d}\bar{z}\ \left(2\bar{\varphi}\bar{u}+\frac{(\partial_{\bar{z}}\bar{u})^2}{2}+\bar{\varphi}^2\right) \\
&= \frac{\bar{a}}{2}\int_0^{\bar{L}} \mathrm{d}\bar{z} \sum_{\bar{q},\bar{q}'\neq 0}\mathrm{e}^{\mathrm{i}(\bar{q}+\bar{q}')\bar{z}}\left(2\bar{\phi}_{\bar{q}}\bar{u}_{\bar{q}'}-\bar{u}_{\bar{q}}\bar{u}_{\bar{q}'}\frac{\bar{q}\bar{q}'}{2}+\bar{\phi}_{\bar{q}}\bar{\phi}_{\bar{q}'}\right) \\
&= \frac{\bar{L}\bar{a}}{2}\sum_{\bar{q}\neq 0}2\bar{\phi}_{\bar{q}}\bar{u}_{-\bar{q}}+\bar{u}_{\bar{q}}\bar{u}_{-\bar{q}}\frac{\bar{q}^2}{2}+\bar{\phi}_{\bar{q}}\bar{\phi}_{-\bar{q}}\,,
\end{aligned} \tag{S29}$$

and

$$\bar{F}_{\bar{b}} = \frac{\bar{b}}{2}\int_0^{\bar{L}} \mathrm{d}\bar{z}\,(\partial_{\bar{z}}\bar{\varphi})^2 = -\frac{\bar{b}}{2}\int_0^{\bar{L}} \mathrm{d}\bar{z} \sum_{\bar{q},\bar{q}'\neq 0}\bar{\phi}_{\bar{q}}\bar{\phi}_{\bar{q}'}\bar{q}\bar{q}'\mathrm{e}^{\mathrm{i}(\bar{q}+\bar{q}')\bar{z}} = \frac{\bar{L}\bar{b}}{2}\sum_{\bar{q}\neq 0}\bar{q}^2\bar{\phi}_{\bar{q}}\bar{\phi}_{-\bar{q}}\,. \tag{S30}$$

Gathering Eqs. (S27)–(S30) yields

$$\begin{aligned}
\bar{F}_{\phi} &= \frac{\bar{L}}{2}\sum_{\bar{q}\neq 0}\left\{\bar{q}^2\left(\bar{\Lambda}-\bar{\mu}+\frac{\bar{a}}{2}\right)\bar{u}_{\bar{q}}\bar{u}_{-\bar{q}}+\left(\bar{\Lambda}\bar{q}^2-2\bar{\mu}+2\bar{a}\right)\bar{u}_{\bar{q}}\bar{\phi}_{-\bar{q}}+\left(\bar{a}+\bar{b}\bar{q}^2\right)\bar{\phi}_{\bar{q}}\bar{\phi}_{-\bar{q}}\right\} \\
&= \frac{\bar{L}}{4}\sum_{\bar{q}\neq 0}\left\{\bar{q}^2\left(\bar{\Lambda}-\bar{a}\right)\bar{u}_{\bar{q}}\bar{u}_{-\bar{q}}+2\bar{\Lambda}\left(\bar{q}^2-1\right)\bar{u}_{\bar{q}}\bar{\phi}_{-\bar{q}}+2\left(\bar{a}+\bar{b}\bar{q}^2\right)\bar{\phi}_{\bar{q}}\bar{\phi}_{-\bar{q}}\right\}.
\end{aligned} \tag{S31}$$

where we inserted $\bar{\mu} = \bar{a} + \bar{\Lambda}/2$ [Eq. (4) of the main text, nondimensionalised] to go to the second line. Combining Eqs. (S23) and (S31) yields Eq. (11) of the main text.

#### S6. FLAT-MEMBRANE LIMIT OF EQ. (11) OF THE MAIN TEXT

We write out Eq. (11) of the main text and return to dimensional units

$$\bar{F} = \frac{\bar{L}}{4}\sum_{\bar{q}\neq 0}\begin{pmatrix}\bar{u}_{\bar{q}} & \bar{\phi}_{\bar{q}}\end{pmatrix}\begin{pmatrix}2\bar{q}^4+\left(\bar{\Lambda}-\bar{a}+2\bar{\sigma}-1\right)\bar{q}^2+\bar{\Lambda}+\bar{a}-2\bar{\sigma}+3 & \bar{\Lambda}(\bar{q}^2-1)\\ \bar{\Lambda}(\bar{q}^2-1) & 2\bar{a}+2\bar{b}\bar{q}^2\end{pmatrix}\begin{pmatrix}\bar{u}_{-\bar{q}}\\ \bar{\phi}_{-\bar{q}}\end{pmatrix}$$

$$\frac{F}{2\pi\kappa} = \frac{L}{4R}\sum_{\bar{q}\neq 0}\begin{pmatrix}\frac{u_q}{R} & \frac{\phi_q}{\Phi}\end{pmatrix}$$

$$\times\begin{pmatrix}2q^4R^4+\left(\frac{\Lambda\Phi R}{\kappa}-\frac{a\Phi^2R^2}{\kappa}+2\frac{\sigma R^2}{\kappa}-1\right)q^2R^2+\frac{\Lambda\Phi R}{\kappa}+\frac{a\Phi^2R^2}{\kappa}-2\frac{\sigma R^2}{\kappa}+3 & \frac{\Lambda\Phi R}{\kappa}\left(q^2R^2-1\right)\\ \frac{\Lambda\Phi R}{\kappa}\left(q^2R^2-1\right) & 2\frac{a\Phi^2R^2}{\kappa}+2\frac{b\Phi^2}{\kappa}q^2R^2\end{pmatrix}\begin{pmatrix}u_{-q}/R\\ \phi_{-q}/\Phi\end{pmatrix}$$

$$F = \frac{\pi L}{2R}\sum_{q\neq 0}\begin{pmatrix}u_q & \phi_q\end{pmatrix}\begin{pmatrix}2\kappa q^4R^2+\left(\Lambda\Phi R-a\Phi^2R^2+2\sigma R^2-\kappa\right)q^2+\frac{\Lambda\Phi}{R}+a\Phi^2-2\sigma+\frac{3\kappa}{R^2} & \Lambda\left(q^2R^2-1\right)\\ \Lambda\left(q^2R^2-1\right) & 2aR^2+2bq^2R^2\end{pmatrix}\begin{pmatrix}u_{-q}\\ \phi_{-q}\end{pmatrix}. \tag{S32}$$

We now find that the leading-order term in the limit of a large radius $R$ reads

$$F = \frac{2\pi RL}{2}\sum_{q\neq 0}\begin{pmatrix}u_q & \phi_q\end{pmatrix}\begin{pmatrix}\kappa q^4+\left(\sigma-a\Phi^2/2\right)q^2 & \frac{1}{2}\Lambda q^2\\ \frac{1}{2}\Lambda q^2 & a+bq^2\end{pmatrix}\begin{pmatrix}u_{-q}\\ \phi_{-q}\end{pmatrix}+\mathcal{O}(1). \tag{S33}$$

The prefactor $2\pi RL$ being the area of the (large-radius) cylinder, we recognise that the above expression is the same free energy per unit area as Eq. (S7).

#### S7. ANALYSIS OF DYNAMICAL INSTABILITY

##### A. General dynamical equations in geometric form

We consider a membrane $\mathcal{M}\subset\mathbb{R}^3$, locally isomorphic to $\mathbb{R}^2$, whose tangent bundle and normal bundle are spanned by a local orthonormal triad, $\{\mathbf{e}_1,\mathbf{e}_2,\mathbf{N}\}$, where $\mathbf{N}$ is normal to the surface and $\mathbf{e}_{1,2}$ are the tangent basis. The surface embedding is defined by the Gauss-Codazzi relations and the shape operator $\mathbf{S}=-\nabla\mathbf{N}$, where $\nabla$ is the gradient operator on the membrane.

We can calculate the elastic forces from the following free energy

$$\mathcal{F} = \int f\,\mathrm{d}A = \int\left(2\kappa H^2-\Lambda\phi H+\frac{a}{2}\phi^2+\frac{b}{2}|\nabla\phi|^2\right)\mathrm{d}A, \tag{S34}$$

where $H=\mathrm{Tr}\,(\mathbf{S})$ is the mean curvature and $\Lambda$, $\kappa$, $a$, and $b$ are as defined in Eqs. (1) and (2) of the main text.. Note that we omit the surface tension, pressure, and chemical potential terms, which will be introduced as conjugate variables to the dynamical constraints. The Gaussian curvature is defined by $K=\det(\mathbf{S})$.

Taking variations of Eq. (S34) with respect to perturbations of the surface of the form $\mathbf{R}\to\mathbf{R}+\psi\mathbf{N}+\delta\mathbf{x}$, where $\delta\mathbf{x}\in T(\mathcal{M})$, we find

$$\begin{aligned}\delta\mathcal{F} &= \int\delta f\,\mathrm{d}A+\int f\,\delta\mathrm{d}A\\ &= \int\left[4\kappa\left(\frac{1}{2}\nabla^2\psi+\psi\left(2H^2-K\right)\right)H-\frac{1}{2}\Lambda\phi\nabla^2\psi-\psi\Lambda\phi(2H^2-K)+b\nabla\phi\cdot\left(\mathbf{N}(\nabla\psi)\cdot\nabla\phi-\psi\nabla\mathbf{N}\cdot\nabla\phi\right)\right]\mathrm{d}A\\ &\quad-\int 2H\left(2\kappa H^2-\Lambda\phi H+\frac{a}{2}\phi^2+\frac{b}{2}|\nabla\phi|^2\right)\mathrm{d}A-\int\left[-\Lambda H+a\phi+b\nabla^2\phi\right]\nabla\phi\cdot\delta\mathbf{x}\,\mathrm{d}A.\end{aligned} \tag{S35}$$

Upon integration by parts, this gives a normal force

$$\begin{aligned}\mathbf{f}^{\mathrm{elastic}} = &-\left(2\kappa\nabla^2H+4\kappa H(H^2-K)-\frac{\Lambda}{2}\nabla^2\phi+\Lambda\phi K-b\nabla\phi\cdot\nabla\mathbf{N}\cdot\nabla\phi-a\phi^2H-b|\nabla\phi|^2H\right)\mathbf{N}\\ &+\left(a\phi+b\nabla^2\phi-\Lambda H\right)\nabla\phi,\end{aligned} \tag{S36}$$

the first part of which is the standard shape equation for an isotropic fluid membrane with purely bending energy [20]. For simplicity, we neglect the tangential elastic forces here.

To derive the full dynamical equations, we will consider a Rayleigh dissipation functional

$$\mathcal{R} = \mathcal{P}^{\text{bulk}} + \mathcal{P}^{\text{mem}} + \mathcal{P}^{\text{constraints}} + \int \frac{\delta \mathcal{F}}{\delta \mathbf{R}} \cdot \mathbf{V} \, \mathrm{d}A \,, \tag{S37}$$

where $\mathbf{V} = \mathrm{d}\mathbf{R}/\,\mathrm{d}t$ is the surface velocity of the membrane. Moreover, $\mathcal{P}^{\text{bulk}}$ is the dissipation functional of the bulk fluid, $\mathcal{P}^{\text{mem}}$ is the dissipation functional of the membrane, and $\mathcal{P}^{\text{constraints}}$ is the dissipation functional associated with constraints on the system dynamics, in our case, fluid and membrane incompressibility and protein conservation.

The bulk dissipation is defined by

$$\mathcal{P}^{\text{bulk}} = \int \eta \mathcal{D} : \mathcal{D} \, \mathrm{d}V \,, \tag{S38}$$

where $\mathcal{D} = \left( \vec{\nabla} \vec{V} + \left( \vec{\nabla} \vec{V} \right)^{\mathrm{T}} \right) /2$ is the stain rate in the ambient fluid (here, $\eta$ is the fluid's shear viscosity). Note that $\vec{\nabla}$ is the gradient operator in $\mathbb{R}^3$ and $\vec{V}$ is the velocity field of the ambient fluid. The associated constraint imposing bulk incompressibility is

$$\mathcal{P}^{\text{bulk.const.}} = \int P \vec{\nabla} \cdot \vec{V} \, \mathrm{d}V, \tag{S39}$$

where $P$ is the hydrodynamic pressure.

Equivalently, for the membrane with surface velocity $\mathbf{V} = \mathbf{v} + v_{\mathrm{n}}\mathbf{N}$, we have

$$\mathcal{P}^{\text{mem}} = \int \eta_m D : D \, \mathrm{d}A \,, \tag{S40}$$

where $\eta_m$ is the 2D membrane viscosity and $D = \mathbb{P} \cdot \left( \nabla \mathbf{V} + (\nabla \mathbf{V})^{\mathrm{T}} \right) \cdot \mathbb{P}/2 = 1/2 \left( \nabla^\alpha v^\beta + \nabla^\beta v^\alpha - S^{\alpha\beta} v_{\mathrm{n}} \right) \mathbf{e}_\alpha \mathbf{e}_\beta$ is the surface deformation rate. Here, $\mathbb{P} = \mathbb{I}_3 - \mathbf{NN}$ is the projection operator onto the tangent space of the membrane from $\mathbb{R}^3$. For the surface incompressibility, we have

$$\mathcal{P}^{\text{mem.const.}} = -\int \sigma \nabla \cdot \mathbf{V} \, \mathrm{d}A = \int \sigma \left( \nabla_\alpha v^\alpha - 2H v_{\mathrm{n}} \right) \mathrm{d}A, \tag{S41}$$

where $\sigma$ is the hydrodynamic surface tension.

Additionally, we impose local conservation of proteins by the dynamical constraint functional

$$\mathcal{P}^{\phi} = \int \mu \left[ \partial_t \phi + \nabla \cdot \left( \mathbf{V}\phi - \mathbf{M} \cdot \nabla \frac{\delta \mathcal{F}}{\delta \phi} \right) \right] \mathrm{d}A \,, \tag{S42}$$

where $\mu$ is the chemical potential of the proteins on the surface and $\mathbf{M}$ is the protein mobility tensor on the surface. For simplicity, we assume $\mathbf{M} = M\mathbb{I}_2$.

Taking functional derivatives of the Rayleigh dissipation functional Eq. (S37) w.r.t. velocity, we find the force balance equations for bulk and the surface, respectively

$$\eta \vec{\nabla}^2 \vec{V} = \vec{\nabla} P, \tag{S43}$$

$$\begin{aligned} &- \left[ 2\kappa \nabla^2 H + 4\kappa H (H^2 - K) - \frac{\Lambda}{2} \nabla^2 \phi + \Lambda \phi K + b \nabla \phi \cdot \mathbf{S} \cdot \nabla \phi - a\phi^2 H - b|\nabla \phi|^2 H \right] \mathbf{N} + \nabla \sigma \\ &+ 2H\sigma \mathbf{N} + \eta_m \Big( \nabla_\| \cdot \left( \nabla_\| \mathbf{v} + (\nabla_\| \mathbf{v})^T \right) - 4 v_n \nabla H - 2\mathbf{S} \cdot \nabla v_n \Big) - \phi \nabla \mu - 2H\mu\phi \mathbf{N} \\ &+ 2\eta_m \big( \nabla_\| \mathbf{v} : (\mathbf{S} - H\mathbb{I}_2) - 2 v_n (H^2 - K) \big) \mathbf{N} = \eta [\mathcal{D}] \cdot \mathbf{N} - [P] \mathbf{N}, \end{aligned} \tag{S44}$$

where $\nabla_\| \mathbf{v} = \nabla \mathbf{v} - (\nabla \mathbf{v} \cdot \mathbf{N})\mathbf{N}$ is the covariant derivative of $\mathbf{v}$.

Functional variation of Eq. (S37) w.r.t. $P$, $\sigma$, and $\mu$ then gives the constraint equations

$$\vec{\nabla} \cdot \vec{V} = 0, \tag{S45a}$$

$$\nabla \cdot \mathbf{V} = 0, \tag{S45b}$$

$$\partial_t \phi + \mathbf{v} \cdot \nabla \phi - aM \nabla^2 \phi + \Lambda M \nabla^2 H + bM \nabla^4 \phi = 0. \tag{S45c}$$

##### B. Linearised axisymmetric equations for a membrane tube

We assume a groundstate with no flow and membrane given by the vector $\mathbf{X}(\theta, z) = ((R+\epsilon u(z))\cos\theta, (R+\epsilon u(z))\sin\theta, z)$, and protein distribution $\phi = \Phi + \epsilon\delta\phi(z)$ where $\epsilon \ll 1$. Moreover, we assume all flows are of order $\epsilon$. We assume forms for the surface tension and chemical potential of the form $\sigma(z) = \sigma_0 + \epsilon\delta\sigma(z)$ and $\mu(z) = \mu_0 + \epsilon\chi\delta\phi$ where $\chi$ is the proteins' 2D inverse compressibility.

Inserting these assumptions into Eqs. (S43)–(S45) gives the following linearised axisymmetric equations for a membrane tube. First, the shape operator amounts to

$$\mathbf{S} = -\nabla\mathbf{N} = -\frac{R-\epsilon u}{R^2}\mathbf{e}_\theta\mathbf{e}_\theta + \epsilon\partial_{zz}u\mathbf{e}_z\mathbf{e}_z, \tag{S46}$$

giving the following mean and Gaussian curvature

$$H = -\frac{1}{2R} + \epsilon\frac{\partial_{zz}u}{2} + \epsilon\frac{u}{2R^2}, \tag{S47a}$$

$$K = -\frac{\epsilon\partial_{zz}u}{R}. \tag{S47b}$$

Second, surface incompressibility [Eq. (S45b)] gives, at first order,

$$\partial_z v^z + \frac{1}{R}\dot{u} = 0. \tag{S48}$$

Third, using Eq. (S44), tangential force balance is given by

$$\partial_z\delta\sigma + 2\eta_m\partial_z^2 v^z - \chi\Phi\partial_z\delta\phi = \eta\vec{e}_z\cdot[\mathcal{D}]\cdot\vec{e}_r, \tag{S49}$$

and normal force balance on the membrane is given by

$$\begin{aligned}
&-\left[\kappa\left(\epsilon\left(\partial_z^4 u + \frac{1}{2R^2}\partial_z^2 u + \frac{3}{2R^4}u\right) - \frac{1}{2R^3}\right) - \frac{\epsilon\Lambda}{2}\partial_z^2\delta\phi - \epsilon\frac{\Lambda\Phi}{R}\partial_z^2 u + \frac{a\Phi^2}{2R} + \epsilon\frac{a\Phi}{R}\delta\phi - \epsilon a\Phi^2\left(\frac{u}{2R^2} + \frac{\partial_z^2 u}{2}\right)\right]\\
&-\frac{\sigma_0}{R} + \epsilon\sigma_0\left(\partial_z^2 u + \frac{u}{R^2}\right) - \epsilon\frac{\delta\sigma}{R} + \frac{\mu_0\Phi}{R} + \epsilon\left(\frac{\mu_0\delta\phi}{R} + \frac{\chi\Phi\delta\phi}{R} - \mu_0\Phi\left(\partial_z^2 u + \frac{u}{R^2}\right)\right) - 2\frac{\eta_m}{R^2}\dot{u}\\
&= \eta\vec{e}_r\cdot[\mathcal{D}]\cdot\vec{e}_r - [P].
\end{aligned} \tag{S50}$$

Fourth, the protein dynamics [Eq. (S45c)] is governed by

$$\partial_t\delta\phi - aM\partial_z^2\delta\phi + \frac{\Lambda M}{2}\left(\partial_z^4 u + \frac{1}{R^2}\partial_z^2 u\right) + bM\partial_z^4\delta\phi = 0. \tag{S51}$$

The solutions to the Stokes equation [Eq. (S43)] in cylindrical coordinates $(r,\theta,z)$ inside $(-)$ and outside $(+)$ the tube are given by

$$\vec{V}^\pm = \vec{\nabla}\zeta^\pm + \vec{\nabla}\times\left(\psi^\pm\vec{e}_z\right) + r\partial_r\vec{\nabla}\xi^\pm + \partial_z\xi^\pm\vec{e}_z\,, \tag{S52a}$$

$$P^\pm = -2\eta\partial_z^2\xi^\pm + P_0^\pm\,, \tag{S52b}$$

$$\left(\zeta^\pm,\psi^\pm,\xi^\pm\right)^{\mathrm{T}} = \int\frac{\mathrm{d}q}{2\pi}\left(Z^\pm,\Psi^\pm,\Xi^\pm\right)^{\mathrm{T}}\Pi_q^\pm(r)e^{iqz}, \tag{S52c}$$

$$\text{where}\quad \Pi_q^\pm(r) = \begin{cases}\Pi_q^+(r) = K_0(qr)\,,\\ \Pi_q^-(r) = I_0(qr)\,,\end{cases}$$

where $I_0(qr)$ and $K_0(qr)$ are modified Bessel functions of the first and second kind.

Taking the Fourier transform in $z$ ($f(z) = \int \frac{dq}{2\pi} \bar{f}_q e^{iqz}$) of the linear perturbations in our Eqs. (S48)–(S51) yields

$$iq\bar{v}_q^z + \frac{1}{R}\partial_t \bar{u}_q = 0, \tag{S53}$$

$$iq\delta\bar{\sigma}_q - 2q^2\eta_m \bar{v}_q^z - iq\chi\Phi\delta\bar{\phi}_q = \mathcal{F}_q\{\eta\vec{e}_z \cdot [\mathcal{D}] \cdot \vec{e}_r\}, \tag{S54}$$

$$\begin{aligned}
&-\Big[\kappa\left(\epsilon\left(q^4 - \frac{1}{2R^2}q^2 + \frac{3}{2R^4}\right)\bar{u}_q - \frac{1}{2R^3}\delta(q)\right) + \frac{\epsilon\Lambda}{2}q^2\delta\bar{\phi}_q + \epsilon\frac{\Lambda\Phi}{R}q^2\bar{u}_q + \frac{a\Phi^2}{2R}\delta(q) + \epsilon\frac{a\Phi}{R}\delta\bar{\phi}_q \\
&- \epsilon a\Phi^2\bar{u}_q\left(\frac{1}{2R^2} - \frac{q^2}{2}\right)\Big] - \frac{\sigma_0}{R}\delta(q) + \epsilon\sigma_0\bar{u}_q\left(\frac{1}{R^2} - q^2\right) - \epsilon\frac{\delta\bar{\sigma}_q}{R} + \frac{\mu_0\Phi}{R}\delta(q) \\
&+ \epsilon\left(\frac{\mu_0\delta\bar{\phi}_q}{R} + \frac{\chi\Phi\delta\bar{\phi}_q}{R} - \mu_0\Phi\bar{u}_q\left(\frac{1}{R^2} - q^2\right)\right) - 2\frac{\eta_m}{R^2}\partial_t\bar{u}_q = \mathcal{F}_q\{\eta\vec{e}_r \cdot [\mathcal{D}] \cdot \vec{e}_r - [P]\}
\end{aligned} \tag{S55}$$

$$\partial_t\delta\bar{\phi}_q = -\left(bMq^4 + aMq^2\right)\delta\bar{\phi}_q + \frac{\Lambda M}{2}\bar{u}_q\left(\frac{q^2}{R^2} - q^4\right). \tag{S56}$$

For $\epsilon = 0$ (the unperturbed state), these are solved for the groundstate conditions found via the free energy analysis, Eqs. (4) and (5) of the main text,

$$\mu_0 = a\Phi + \frac{\Lambda}{2R},$$
$$\frac{P_0 R^3}{\kappa} = \frac{\sigma_0 R^2}{\kappa} - \frac{1}{2} - \frac{a\Phi^2R^2}{2\kappa} - \frac{\Lambda\Phi R}{2\kappa}.$$

We solve the ambient Stokes equations in Fourier space for no-slip boundary conditions. We have $\Psi^{\pm} = 0$ as the system is axisymmetric. The other coefficients are computed as

$$Z^+ = \frac{K_1(qR)(qR\partial_t\bar{u}_q - i\bar{v}_q^z) - K_0(qR)(\partial_t\bar{u}_q + iqR\bar{v}_q^z)}{q\left[qRK_0(qR)^2 + 2K_1(qR)K_0(qR) - qRK_1(qR)^2\right]}, \tag{S57a}$$

$$Z^- = \frac{I_1(qR)(-qR\partial_t\bar{u}_q + i\bar{v}_q^z) - I_0(qR)(\partial_t\bar{u}_q + iqR\bar{v}_q^z)}{q\left[qRI_0(qR)^2 - 2I_1(qR)I_0(qR) - qRI_1(qR)^2\right]}, \tag{S57b}$$

$$\Xi^+ = \frac{\partial_t\bar{u}_qK_0(qR) - i\bar{v}_q^zK_1(qR)}{q\left[qRK_0(qR)^2 + 2K_1(qR)K_0(qR) - qRK_1(qR)^2\right]}, \tag{S57c}$$

$$\Xi^- = \frac{\partial_t\bar{u}_qI_0(qR) + i\bar{v}_q^zI_1(qR)}{q\left[qRI_0(qR)^2 - 2I_1(qR)I_0(qR) - qRI_1(qR)^2\right]}. \tag{S57d}$$

The ambient velocities and pressures in Fourier space are then given by

$$\bar{V}_q^+ = \left[q^2r\Xi^+\left(K_0(qr) + \frac{K_1(qr)}{qr}\right) - qZ^+K_1(qr)\right]\vec{e}_r + \left[iqr\Xi^+\left(\frac{K_0(qr)}{r} - qK_1(qr)\right) + iqZ^+K_0(qr)\right]\vec{e}_z, \tag{S58a}$$

$$\bar{V}_q^- = \left[q^2r\Xi^-\left(I_0(qr) - \frac{I_1(qr)}{qr}\right) + qZ^-I_1(qr)\right]\vec{e}_r + \left[iqr\Xi^-\left(\frac{I_0(qr)}{r} + qI_1(qr)\right) + iqZ^-I_0(qr)\right]\vec{e}_z, \tag{S58b}$$

$$\bar{P}^+ = P_0^+ + 2\eta q^2\Xi^+K_0(qr), \tag{S58c}$$

$$\bar{P}^- = P_0^- + 2\eta q^2\Xi^-I_0(qr). \tag{S58d}$$

The tangential force balance and continuity equations can be solved for the surface tension variation, giving

$$\begin{aligned}
\delta\bar{\sigma}_q =& \delta\bar{\phi}_q\Phi\chi + \frac{2\eta_m\partial_t\bar{u}_q}{R} \\
&+ \frac{2\eta\partial_t\bar{u}_q\left\{qRI_0(qR)\left[qRK_0(qR) + K_1(qR)\right] - I_1(qR)\left[\left(q^2R^2 + 2\right)K_1(qR) + qRK_0(qR)\right]\right\}}{q^2R^2\left[qRI_0(qR)^2 - 2I_1(qR)I_0(qR) - qRI_1(qR)^2\right]\left[qRK_0(qR)^2 + 2K_1(qR)K_0(qR) - qRK_1(qR)^2\right]},
\end{aligned} \tag{S59}$$

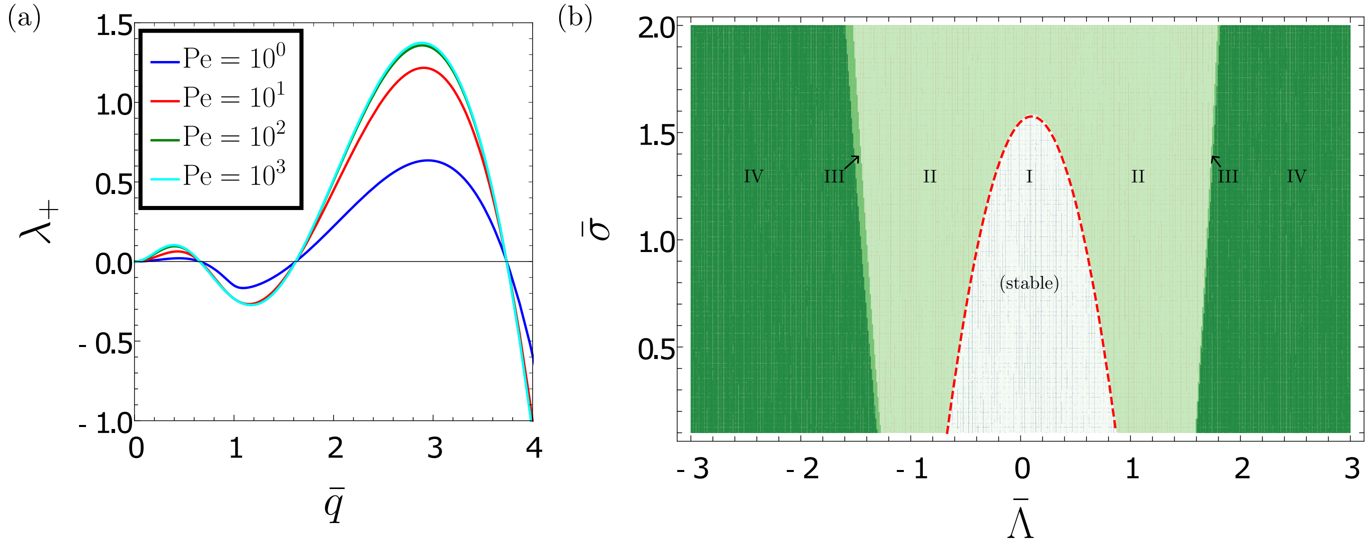


Figure S3. (a) Largest eigenvalues for the dynamical instability for $\bar{\Lambda} = 2$, $\mathrm{Bq} = 1$, and several values of Péclet number, Pe. All other parameters are the same as in Fig. 2 of the main text. (b) Stability plot for $\mathrm{Pe} = 1$; all other values are identical to Fig. 3(c) of the main text. The only effect is the shrinking of region III.

which can then be substituted back into the shape equation, Eq. (S55), we find

$$-\frac{\kappa}{2R^4}\left(2\bar{q}^4 + \zeta_2\bar{q}^2 + \zeta_0\right)\bar{u}_q - \frac{\Lambda}{2R^3}\left(\bar{q}^2 - 1\right)\delta\bar{\phi}_q = B'\partial_t\bar{u}_q, \tag{S60}$$

where

$$\begin{aligned} B' &= \frac{2\eta}{R^2}\left(\frac{\left(\bar{q}^2+1\right)\left[I_0(\bar{q})K_1(\bar{q}) - I_1(\bar{q})K_2(\bar{q})\right]}{\bar{q}\left(\bar{q}I_0(\bar{q})^2 - 2I_1(\bar{q})I_0(\bar{q}) - \bar{q}I_1(\bar{q})^2\right)\left[\bar{q}K_0(\bar{q})^2 + 2K_1(\bar{q})K_0(\bar{q}) - \bar{q}K_1(\bar{q})^2\right]} + 2\frac{\eta_m}{R\eta}\right) \\ &\equiv \frac{2\eta}{R^2}\left(\beta + 2\frac{\eta_m}{R\eta}\right), \end{aligned} \tag{S61}$$

and $\zeta_2 = \bar{\Lambda} - \bar{a} + 2\bar{\sigma} - 1$, $\zeta_0 = \bar{\Lambda} + \bar{a} - 2\bar{\sigma} + 3$ are the same as those defined in the main text. For completeness,

$$\beta = \frac{\left(\bar{q}^2+1\right)\left[I_0(\bar{q})K_1(\bar{q}) - I_1(\bar{q})K_2(\bar{q})\right]}{\bar{q}\left(\bar{q}I_0(\bar{q})^2 - 2I_1(\bar{q})I_0(\bar{q}) - \bar{q}I_1(\bar{q})^2\right)\left[\bar{q}K_0(\bar{q})^2 + 2K_1(\bar{q})K_0(\bar{q}) - \bar{q}K_1(\bar{q})^2\right]}. \tag{S62}$$

With the Boussinesq number $\mathrm{Bq} = \eta_m/R\eta$, Péclet number $\mathrm{Pe} = M\eta/R^3\Phi^2$, and nondimensionalisation with respect to the timescale $\eta R/\kappa$, lengthscale $R$, and concentration $\Phi$, we find Eq. (16) of the main text,

$$\partial_t\begin{pmatrix}\bar{u}_q \\ \delta\bar{\phi}_q\end{pmatrix} = \begin{pmatrix}\frac{-1}{4(\beta+2\mathrm{Bq})}\left(2\bar{q}^4 + \zeta_2\bar{q}^2 + \zeta_0\right) & \frac{-\bar{\Lambda}}{4(\beta+2\mathrm{Bq})}\left(\bar{q}^2 - 1\right) \\ -\frac{\mathrm{Pe}}{2}\bar{\Lambda}\left(\bar{q}^4 - \bar{q}^2\right) & -\mathrm{Pe}\left(\bar{a}\bar{q}^2 + \bar{b}\bar{q}^4\right)\end{pmatrix}\begin{pmatrix}\bar{u}_q \\ \delta\bar{\phi}_q\end{pmatrix} \tag{S63}$$

where $\bar{a}$, $\bar{b}$, and $\bar{\Lambda}$ are as defined in the main text. Note that this matrix is precisely $B$ [Eq. (11) of the main text], with its elements divided by various dissipative factors and an overall minus sign. The eigenvalues of the matrix (*i.e.* the growth rate) are plotted in Fig. 3 of the main text. In addition, Fig. S3(a) shows that the growth saturates for large Péclet number. Figure S3(b) shows a stability diagram for a smaller value of Péclet number than used in the main text.

#### S8. ENERGY-MINIMISING SHAPES

To determine the energy-minimising shapes, we start by rewriting the dimensionless free energy $\bar{F}$ of a membrane tube coated by proteins as

$$2\pi\bar{F} = \int \mathrm{d}\bar{A}\left(2\left(\bar{H} - \bar{C}(\bar{\phi})\right)^2 + \tilde{\sigma}(\bar{\phi}) + R^2\frac{\bar{b}}{2}|\nabla\bar{\phi}|^2\right) - \bar{P}\int \mathrm{d}\bar{V} \tag{S64}$$

with $\mathrm{d}\bar{A} = R^{-2}\,\mathrm{d}A$, $\mathrm{d}\bar{V} = R^{-3}\,\mathrm{d}V$, $\bar{C}(\bar{\phi}) = \frac{\bar{\Lambda}}{4}\bar{\phi}$, and $\widetilde{\sigma}(\bar{\phi}) = \bar{\sigma} - \left(\bar{a} + \frac{\bar{\Lambda}}{2}\right)\bar{\phi} + \left(\frac{\bar{a}}{2} - \frac{\bar{\Lambda}}{8}\right)\bar{\phi}^2$. At the boundary between a dilute and a dense protein domain the term $R^2\frac{\bar{b}}{2}|\nabla\bar{\phi}|^2$ acts a line tension. To estimate the magnitude of the line tension, we consider a cylindrical tube with dimensionless radius $\bar{r}_\mathrm{b}$. The protein density shall change over a dimensionless length $\bar{l}$ from a dense domain with density $\bar{\phi}^{(I)}$ to a dilute domain, with $\bar{\phi}^{(II)}$. If the mean curvature follows the spontaneous curvature ($\bar{H} = \bar{C}$), the contribution of the domain boundary to the free energy $\bar{F}_\mathrm{b}$ is thus approximated by

$$2\pi\bar{F}_\mathrm{b} = 2\pi\bar{r}_\mathrm{b}\bar{l}\left(\bar{\sigma} + \frac{\bar{b}}{2}\left(\frac{\bar{\phi}^{(I)} - \bar{\phi}^{(II)}}{\bar{l}}\right)^2\right), \tag{S65}$$

where we use $\widetilde{\sigma} \approx \bar{\sigma}$. Minimising $\bar{F}_\mathrm{b}$ with respect to $\bar{l}$ and setting $\bar{l}$ back into Eq. (S65) leads to

$$2\pi\bar{F}_\mathrm{b} = 2\pi\bar{r}_\mathrm{b}\gamma, \qquad \text{with} \qquad \gamma = \sqrt{2\bar{b}\bar{\sigma}}\left(\bar{\phi}^{(I)} - \bar{\phi}^{(II)}\right). \tag{S66}$$

The free energy of the coated membrane tube is now written as

$$2\pi\bar{F}_\mathrm{b} = \widetilde{F} + 2\pi\bar{r}_\mathrm{b}\gamma, \qquad \text{with} \qquad \widetilde{F} = \int \mathrm{d}\bar{A}\left(\left(\bar{H} - \bar{C}(\bar{\phi})\right)^2 + \widetilde{\sigma}(\bar{\phi})\right) - \bar{P}\int \mathrm{d}\bar{V}\,. \tag{S67}$$

Within a domain with constant $\bar{\phi}$, the functional variation of $\widetilde{F}$ leads to the following shape equation [7]:

$$\bar{P} = -2\widetilde{\sigma}(\bar{\phi})\bar{H} + 2\left[\nabla^2\bar{H} + 2\left(\bar{H} - \bar{C}(\bar{\phi})\right)\left(\bar{H}^2 + \bar{H}\bar{C}(\bar{\phi}) - \bar{K}_\mathrm{g}\right)\right], \tag{S68}$$

with $\bar{K}_\mathrm{g}$ the dimensionless Gaussian curvature. A shape with constant mean curvature is a solution to Eq. (S68), if

$$\bar{P} = -2\widetilde{\sigma}(\bar{\phi})\bar{H}\,, \tag{S69a}$$

$$2\left(\bar{H} - \bar{C}(\bar{\phi})\right)\left(\bar{H}^2 + \bar{H}\bar{C}(\bar{\phi}) - \bar{K}_\mathrm{g}\right) = 0\,. \tag{S69b}$$

In the following, we discuss the solutions of Eq. (S68) for different $\bar{\Lambda}$.

1. **negative $\bar{\Lambda}$, one continuous protein domain:**
The protein density $\bar{\phi}$ sets the spontaneous curvature $\bar{C}$. To fulfil the conditions in Eq. (S69), the mean curvature has to equal the spontaneous curvature $\bar{H} = \bar{C}(\bar{\phi})$. To fully describe the shape, we must specify the mean curvature and the minimum radius $\bar{r}_\mathrm{min}$. For a given $\bar{\phi}$, $\bar{r}_\mathrm{min}$ is varied so that the volume of the undulating shape is equal to the volume of a cylinder of the same length and with radius $\bar{r} = 1$. Subsequently, $\bar{\phi}$ is varied to determine the shape with minimal energy.
For $\bar{\Lambda} = -1.8$, we find $\bar{\phi} = 1.112$, $\bar{H} = -0.5004$ and $\bar{r}_\mathrm{min} = 0.96$.

2. **positive $\bar{\Lambda}$, one continuous protein domain:**
In analogy to the case for negative $\bar{\Lambda}$, we find that a solution to the shape equation has to fulfil $\bar{H} = \bar{C}(\bar{\phi})$ and thus $\bar{H} = \bar{\Lambda}\bar{\phi}/4$. Since both $\bar{\Lambda}$ and $\bar{\phi}$ are positive, $\bar{H}$ has to be positive as well. There are no cylindrically symmetric, non-self-intersecting shapes with $\bar{H} > 0$. Hence, there is no physically meaningful solution to the shape equation.

3. **negative $\bar{\Lambda}$, alternation of dense and dilute domains:**
The proteins densities $\bar{\phi}^{(I)}$ (dense) and $\bar{\phi}^{(II)}$ (dilute) set a spontaneous curvature in the respective domain. Condition Eq. (S69b) is fulfilled if the mean curvatures in each domain equal the spontaneous curvature. Furthermore, according to condition Eq. (S69a) the two densities are linked via

$$\widetilde{\sigma}(\bar{\phi}^{(I)})\bar{C}(\bar{\phi}^{(I)}) = \widetilde{\sigma}(\bar{\phi}^{(II)})\bar{C}(\bar{\phi}^{(II)}), \tag{S70}$$

since the pressure does not vary along the tube. For a given $\bar{\phi}^{(I)}$, the corresponding $\bar{\phi}^{(II)}$ and thus mean curvatures in the two domains are set. To fully describe the tube shape, we need to find the minimal tube radius $\bar{r}_\mathrm{min}$ and the radius at the boundary of the dense and dilute domain $\bar{r}_\mathrm{b}$. To determine the two parameters, we start by setting $\bar{\phi}^{(I)}$ and $\bar{r}_\mathrm{min}$ to fixed values and adjust $\bar{r}_\mathrm{b}$ such that volume conservation is achieved. Subsequently, we vary $\bar{\phi}^{(I)}$ and $\bar{r}_\mathrm{min}$ to find the shape with the lowest energy.
For $\bar{\Lambda} = -1.8$, we find $\bar{\phi}^{(I)} = 2.763$, which corresponds to $\bar{\phi}^{(II)} = 0$, $\bar{H}^{(I)} = -1.243$, $\bar{H}^{(II)} = 0$ and $\bar{r}_\mathrm{min} = 0.7$, $\bar{r}_\mathrm{b} = 1.322$.

4. **positive $\bar{\Lambda}$, alternation of dense and dilute domains:**
We again note that the protein densities set the spontaneous curvature in the dense and dilute domains. However, as discussed under point 2, a physically meaningful shape cannot have a positive mean curvature everywhere. Instead, a feasible solution must exhibit alternations between positive and negative mean curvature domains. We note that, for $\bar{C} = 0$, *i.e.* in a protein-free domain, condition Eq. (S69b) is fulfilled if the square of the mean curvature and the Gaussian curvature are identical. Or, equivalently, the two principle curvatures have to be identical. A shape with two identical, negative principle curvatures corresponds to a segment of a sphere.
Based on these notions, we look for a shape that alternates between a protein-free domain $\bar{\phi}^{(II)} = 0$, with a negative mean curvature $\bar{H}^{(II)}$ that follows a spherical shape and a protein-coated domain $\bar{\phi}^{(I)}$, where the mean curvature follows the spontaneous curvature, $\bar{H}^{(II)} = \bar{C}(\bar{\phi}^{(I)})$. The protein density $\bar{\phi}^{(I)}$ is set by Eq. (S69a):
$$\widetilde{\sigma}(\bar{\phi}^{(I)})\bar{C}(\bar{\phi}^{(I)}) = \widetilde{\sigma}(0)\bar{H}^{(II)}. \tag{S71}$$
To fully describe the tube shape, we need to find the minimal tube radius $\bar{r}_{\mathrm{min}}$ and the radius at the boundary of the dense and dilute domain $\bar{r}_{\mathrm{b}}$. We follow a similar procedure as described in point 3. For a fixed $\bar{r}_{\mathrm{min}}$, $\bar{r}_{\mathrm{b}}$ is adjusted to ensure volume conservation. Subsequently, $\bar{r}_{\mathrm{min}}$ is varied to find the energy-minimising shape. For $\bar{\Lambda} = 1.8$, we find $\bar{\phi}^{(I)} = 1.002$, and $\bar{r}_{\mathrm{min}} = 0.291$, $\bar{r}_{\mathrm{b}} = 0.29$, which implies $\bar{H}^{(I)} = 0.451$ and $\bar{H}^{(II)} = -0.772$.

---
